# Metal cofactor stabilization by a partner protein is a widespread strategy employed for amidase activation

**DOI:** 10.1101/2022.01.21.477235

**Authors:** Julia E. Page, Meredith A. Skiba, Truc Do, Andrew C. Kruse, Suzanne Walker

## Abstract

Construction and remodeling of the bacterial peptidoglycan (PG) cell wall must be carefully coordinated with cell growth and division. Central to cell wall construction are hydrolases that cleave bonds in peptidoglycan. These enzymes also represent potential new antibiotic targets. One such hydrolase, the amidase LytH in *Staphylococcus aureus*, acts to remove stem peptides from PG, controlling where substrates are available for insertion of new PG strands and consequently regulating cell size. When it is absent, cells grow excessively large and have division defects. For activity, LytH requires a protein partner, ActH, that consists of an intracellular domain, a large rhomboid protease domain, and three extracellular tetratricopeptide repeats (TPRs). Here we demonstrate that the amidase-activating function of ActH is entirely contained in its extracellular TPRs. We show that ActH binding stabilizes metals in the LytH active site, and that LytH metal binding in turn is needed for stable complexation with ActH. We further present a structure of a complex of the extracellular domains of LytH and ActH. Our findings suggest that metal cofactor stabilization is a general strategy used by amidase activators and that ActH houses multiple functions within a single protein.

**SIGNIFICANCE STATEMENT:** The Gram-positive pathogen *Staphylococcus aureus* is a leading cause of antibiotic resistance-associated death in the United States. Many antibiotics used to treat *S. aureus*, including the beta-lactams, target biogenesis of the essential peptidoglycan (PG) cell wall. Some hydrolases play important roles in cell wall construction and are potential antibiotic targets. The amidase LytH, which requires a protein partner, ActH, for activity, is one such hydrolase. Here, we uncover how the extracellular domain of ActH binds to LytH to stabilize metals in the active site for catalysis. This work advances our understanding of how hydrolase activity is controlled to contribute productively to cell wall synthesis.

## INTRODUCTION

The peptidoglycan cell wall is an essential component of the cell envelope that maintains cell integrity, size, and morphology (1). Its building block, Lipid II, is synthesized inside the cell, flipped across the cell membrane, and then polymerized and crosslinked by peptidoglycan synthases from the penicillin binding protein and SEDS (shape, elongation, division and sporulation) families (2, 3). These enzymes have received much attention, particularly because the penicillin binding proteins are the target of penicillin and other beta-lactams, one of the most successful classes of antibiotics in the clinic (4). However, many other enzymes, including hydrolases, are integral to building mature cell wall (5). Hydrolases are diverse enzymes that cleave bonds in peptidoglycan to allow growth, cell separation, cell wall recycling, and more (5, 6). These enzymes play important roles in bacterial physiology and present novel opportunities for antibiotic development, particularly for use in combination with beta-lactams (7, 8, 9).

LytH, a membrane-bound amidase from *Staphylococcus aureus* that acts early in cell wall synthesis, is important in controlling cell growth and division (7). It removes stem peptides from the glycan backbone of membrane-proximal peptidoglycan to control the availability of substrates for insertion of new strands (Fig. 1). When LytH is absent, cells grow excessively large and have misplaced division septa. LytH mutants also display increased sensitivity to beta-lactams. Because excessive cell wall cleavage can lead to lysis, hydrolases must be carefully controlled. For activity, LytH requires another membrane protein called ActH. Knockouts of ActH share phenotypes of Δ*lytH* mutants, including cell size and division defects as well as oxacillin sensitivity (7). Activators of other cell wall hydrolases have been identified (10–15), but ActH does not share homology with any of them. ActH therefore provides a new opportunity to learn how amidase activity is controlled in Gram-positive organisms.

**Fig. 1.**
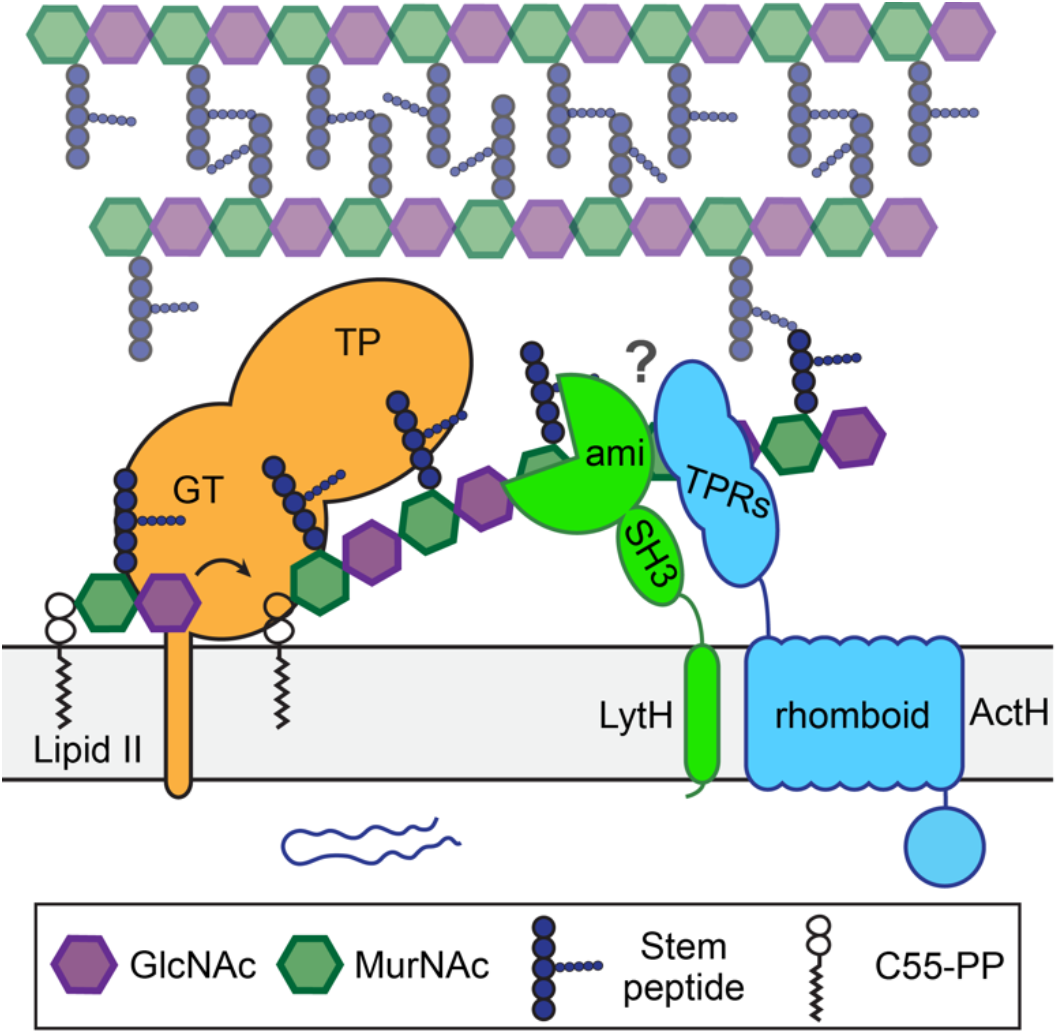
How ActH stimulates the amidase activity of LytH is unknown. Glycosyltransferases (GTs) polymerize Lipid II into glycan strands that get crosslinked into the cell wall by transpeptidases (TPs). LytH-ActH cleaves stem peptides off of uncrosslinked nascent peptidoglycan, controlling the availability of stem peptides that can be used as transpeptidation substrates for insertion of new peptidoglycan strands. ActH is required for robust amidase activity of LytH, but how the two proteins interact to produce this activity is unknown. LytH contains a TM helix, an SH3 domain, and a catalytic amidase_3 domain (ami). ActH has a predicted intracellular domain of 150 amino acids, a rhomboid protease domain, and an extracellular domain with three tetratricopeptide repeats (TPRs).

In this work, we combine structural studies with biochemical and cellular experiments to elucidate how LytH and ActH interact to produce amidase activity. Beyond advancing our understanding of the LytH-ActH complex, which serves as a potential target for beta lactam potentiators, this work reveals principles that likely extend to other hydrolase activators beyond *S. aureus*.

## RESULTS

### The LytH amidase domain and ActH TPRs are sufficient for amidase activity *in vitro*

We first sought to determine what portions of LytH and ActH are necessary to produce amidase activity. ActH is predicted to contain a cytoplasmic domain, a 7 transmembrane helix (TM) domain with homology to the rhomboid proteases, and an extracellular domain with three tetratricopeptide repeats (TPRs). LytH contains a single transmembrane helix, an SH3 (Src homology 3) domain, and a zinc-dependent amidase domain (Fig. 1, 2A). To identify which domains of these proteins are required for amidase activity, we polymerized fluorophore-labeled Lipid II, treated the peptidoglycan oligomers with pairs of truncated or full-length LytH and ActH proteins, and analyzed the products by SDS-PAGE (Fig. 2B). Because the oligomers are labeled on the stem peptide, amidase activity produces tighter spacing of the peptidoglycan ladder with loss of signal intensity; a new band representing the released stem peptide also appears in the middle of the gel. Full-length LytH on its own has a small amount of activity, evidenced by some lightening of the peptidoglycan oligomer ladder and a faint band for the released stem peptide after five hours. When ActH is added, LytH activity increases substantially (compare Fig. 2C lanes 4 and 7), producing a strong signal for the released stem peptide and new, low molecular weight ladder bands. Like full-length LytH, LytH constructs lacking either just the TM helix (LytH_ΔTM_) or both the TM helix and the SH3 domain (LytH_ami_) have minimal activity in the absence of ActH. However, when combined with ActH, they produce full amidase activity (Fig. 2C), showing that the LytH amidase domain does not require the SH3 domain or TM helix for activity or to be activated.

**Fig. 2.**
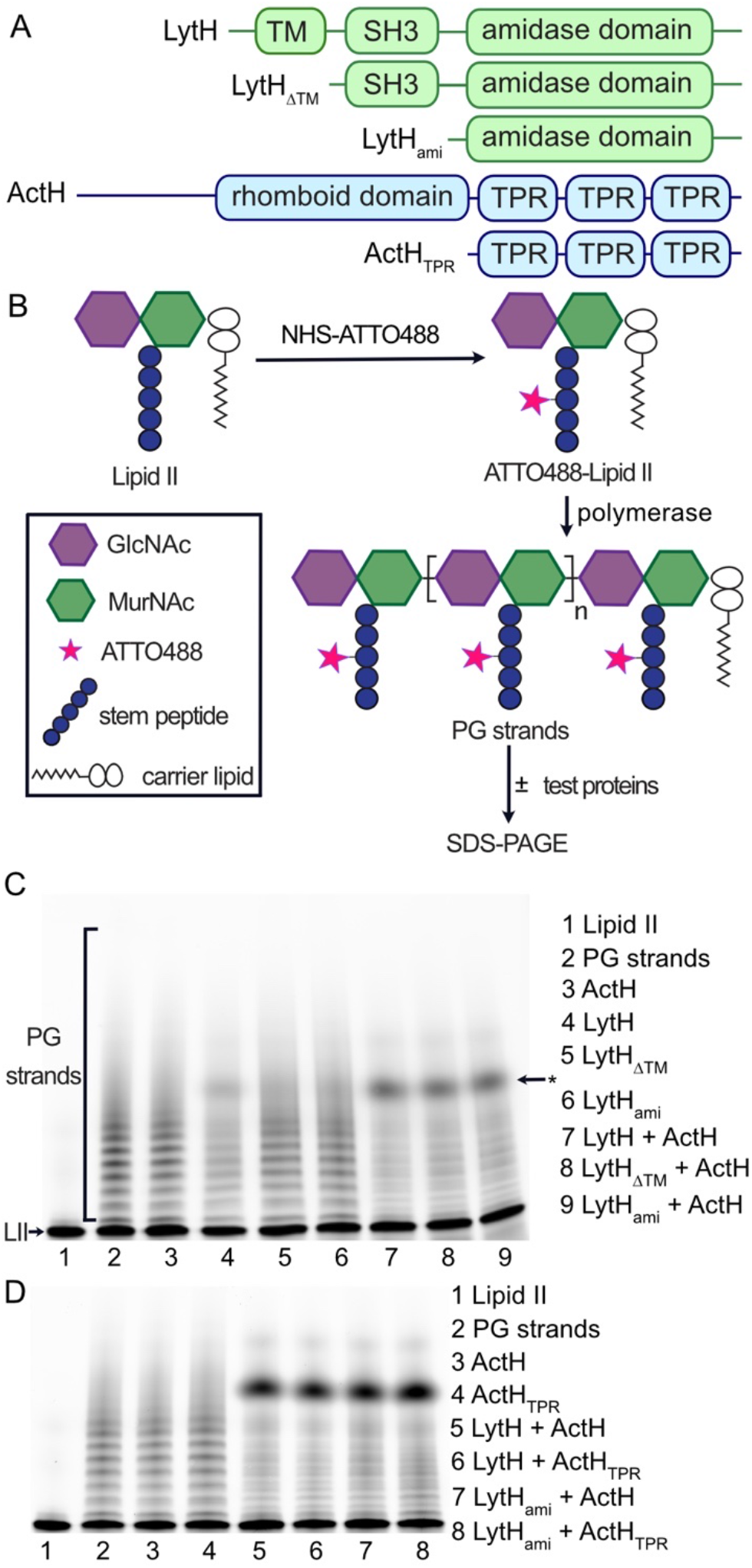
The LytH amidase domain and ActH TPRs are sufficient for amidase activity *in vitro*. **(*A*)** Domain structure of LytH and ActH and truncation mutants tested for activity. (*B*) To detect amidase activity, fluorophore-labeled Lipid II is polymerized into uncrosslinked peptidoglycan strands, treated with the enzyme of interest, and visualized by SDS-PAGE and fluorescence imaging. (*C*) The LytH amidase domain alone (LytH_ami_, LytH[102-291]) has minimal activity but can be activated by ActH. * indicates the released fluorophore-labeled stem peptide. (*D*) The ActH TPRs (ActH_TPR_, ActH[365-487]) are sufficient to activate LytH.

**Fig. 3.**
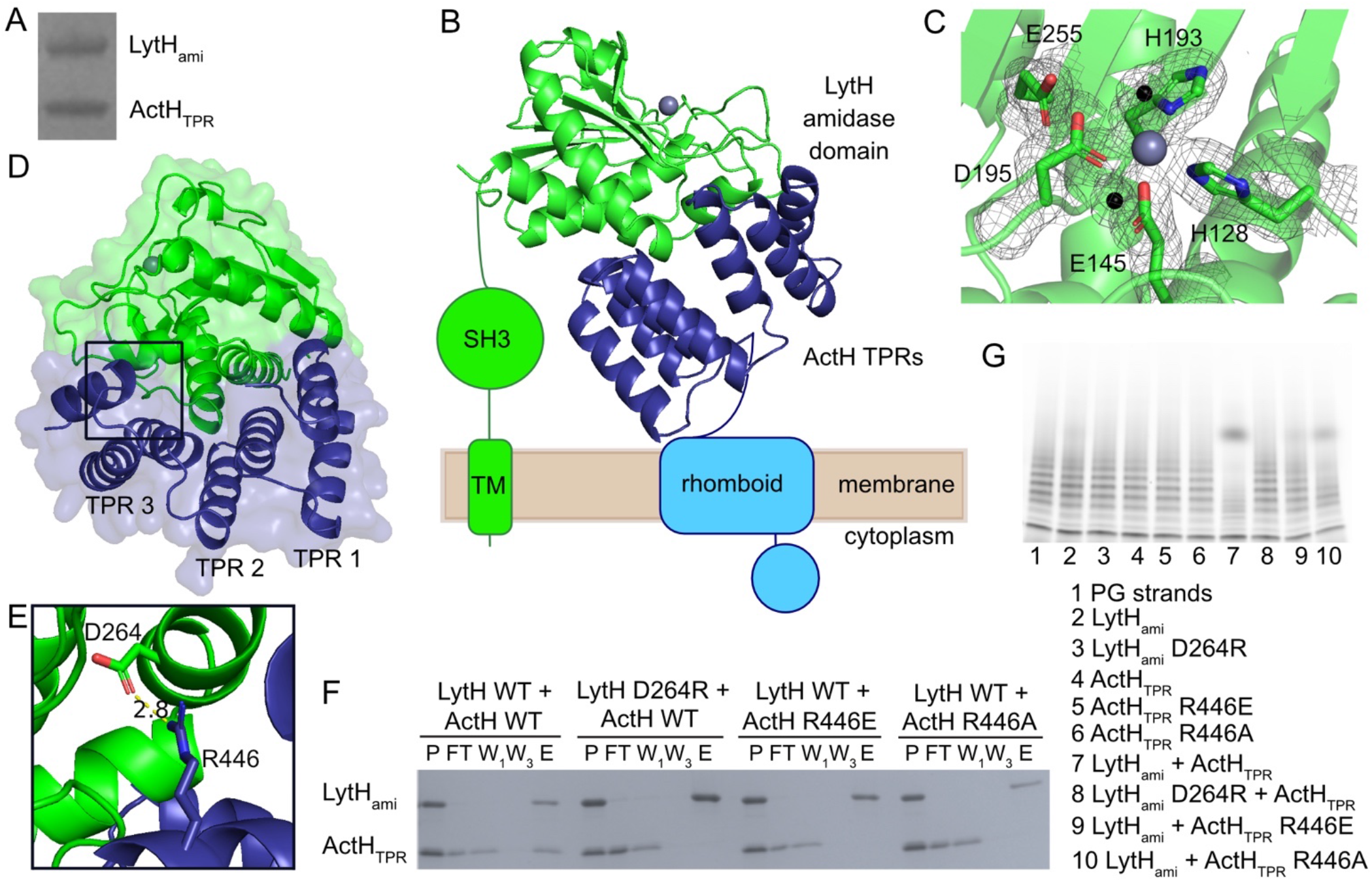
Structure of the LytH amidase domain in complex with the ActH TPRs shows an extensive interface with contacts from LytH alpha helices to the TPRs. (*A*) The LytH amidase domain (LytH[117-291]) and ActH TPRs (ActH[365-479]) co-purify as a stable 1:1 complex from *E. coli*. (*B*) A crystal structure of LytH[117-291, R245A]-ActH[365-479] shows the LytH amidase domain atop the ActH TPRs. The rest of LytH and ActH are schematized. (*C*) Four amino acid side chains (H128, E145, H193, and D195) and two waters (black spheres) coordinate zinc in the active site of LytH. E255 is conserved throughout amidase_3 family proteins. 2Fo-Fc electron density (gray mesh) is contoured at 1σ. (*D*) There is an extensive interface between alpha helices of the LytH amidase domain and the concave surface of the ActH TPRs. (*E*) A salt bridge forms between LytH D264 and ActH R446. (*F*) FLAG-tagged LytH_ami_ (LytH[102-291]) bound to α-FLAG resin pulls down wild-type ActH_TPR_ (ActH[365-487]). LytH_ami_ D264R, ActH_TPR_ R446E, and ActH_TPR_ R446A are no longer able to stably form a complex with their wild-type partner protein. For each sample, P = pre-loading, FT = FLAG resin flow-through, W1 = wash 1, W3 = wash 3, and E = elution. WT = wild-type. (*G*) Fluorescently labeled PG oligos were treated with individual proteins or combinations of LytH_ami_ and ActH_TPR_. Mutants in ActH and LytH that disrupt complex binding correspondingly diminish amidase activity.

We next wondered what portions of ActH are needed to stimulate LytH activity. Given that the ActH TPRs are located extracellularly in proximity to the LytH amidase domain and that TPRs are known to mediate protein-protein interactions (16), we posited that the TPRs of ActH might be responsible for its activation of LytH. When combined with either full-length LytH or the LytH amidase domain alone, the ActH TPRs (ActH_TPR_) stimulated amidase activity equivalently to full-length ActH (Fig. 2D). The extracellular components of LytH and ActH are therefore sufficient for amidase activity *in vitro*.

### The LytH amidase domain and ActH TPRs have an extensive binding interface

To understand the molecular interactions between LytH and ActH, we desired to crystallize LytH-ActH but were unsuccessful in obtaining a structure of the full-length membrane protein complex. Knowing that the soluble domains are sufficient for activity, we wondered if they might also form a stable complex that could be crystallized. We found that LytH_ami_ and ActH_TPR_ co-purified from *E. coli* as a stable 1:1 complex (Fig. 3A). We were able to crystallize this complex, but pathologies in the crystal lattice impeded refining the structure. We substituted a single amino acid in LytH_ami_ to disrupt a lattice contact and were able to solve the structure of that complex to 1.8 Å resolution (Fig. 3B, 3C, Supplementary Table 1).

The structure shows the ActH TPRs binding to the base of the LytH catalytic domain on the opposite side from the active site. Each of the three TPRs demonstrates the classic helical hairpin of these structural elements (16). The TPRs are connected by short loops, forming a halfpipe with concave and convex surfaces. TPRs most commonly bind unstructured peptides in an extended conformation along the concave surface (17). The concave surface of the ActH TPR domain interacts with LytH; however, the bound region of LytH is structured and alpha-helical (Fig. 3D). The ActH TPRs are one of only a handful of TPR domains known to bind globular proteins (17–19).

By solvent accessibility analysis (20), LytH and ActH have a large interface with an area of 1024 Å^2^ stabilized by twelve hydrogen bonds and two salt bridges. One of these salt bridges is between LytH D264 and ActH R446 (Fig. 3E). To test the importance of the LytH-ActH interface observed in the crystal for protein complex formation, we used a simple *in vitro* pull-down experiment to test mutants disrupting this salt bridge. When we mixed FLAG-tagged LytH_ami_ and His-tagged ActH_TPR_, incubated them with FLAG resin, washed, and then eluted with FLAG peptide, both proteins were seen in the elution in approximately a 1:1 ratio. However, when LytH_ami_ D264R, ActH_TPR_ R446E, or ActH_TPR_ R446A was combined with the wild-type version of its respective partner, no ActH_TPR_ was observed in the elution (Fig. 3F). This salt bridge is thus an important point of contact in the LytH-ActH binding interface, supporting the functional relevance of the binding orientation between ActH and LytH observed in the crystal structure. This binding is also important for amidase activity. LytH_ami_ D264R had no activity with or without ActH_TPR_. ActH_TPR_ R446E was also unable to activate LytH_ami_. ActH_TPR_ R446A modestly activated LytH_ami_ (Fig. 3G), suggesting that this mutant retains some ability to bind to LytH_ami_, although the interaction was too weak to observe in the pull-down. The more dramatic effect of the LytH_ami_ D264R and ActH_TPR_ R446E substitutions is consistent with the direct charge-charge repulsions created by those mutations.

### LytH has four amino acids coordinating zinc, but one is dispensable for zinc binding

The LytH amidase domain consists of a twisted six-stranded beta sheet surrounded by six alpha helices. The fold is highly conserved with solved structures of other proteins in the amidase_3 family (21–30; PDB 1JWQ, 3CZX, and 4RN7). In our crystal structure, we observed a metal ion in the active site (initially assumed to be zinc, but see below) with an octahedral coordination sphere made up of four amino acid side chains (H128, E145, H193, D195) and two water molecules (Fig. 3C). The coordinating histidines and glutamate are conserved across this amidase family, but D195 is not strictly conserved (Fig. S1A). In many other amidases, this aspartate is an asparagine that is flipped out toward the solvent (Fig. S1B). In these amidases, only three amino acid side chains (corresponding to H128, E145, and H193) coordinate zinc, and the remaining ligands are water molecules. Like LytH, *E. coli* AmiB and AmiC, which also require protein activators, have an aspartate that is positioned similarly to D195 to coordinate zinc (22, 23) (Fig. S1C). In AmiB and AmiC, an alpha helix blocks the active site, and the activators are presumed to cause a conformational change that exposes the active site. In LytH, the active site is already exposed on the surface of the protein, raising the question of why it is inactive without ActH (Fig. S2).

Our previous studies have shown that LytH D195 is required for catalytic activity (7). Because it is not strictly conserved, we wanted to test if it is also required for zinc binding. We co-purified ActH_TPR_ with wild-type or D195A LytH_ami_, as well as with mutants of two of the three other zinc-coordinating residues, H128A and E145A, in buffer without added metal ions. We then used inductively coupled plasma mass spectrometry (ICP-MS) to measure levels of zinc and several other transition metals in these proteins. The wild-type sample contained predominantly zinc and iron in similar amounts, adding up to about 0.4 equivalents of metal per LytH complex. Although LytH is known as a zinc-dependent amidase, we wondered if iron is also bound in the LytH active site. To determine the metals present at the metal binding site in our crystal structure, we used anomalous scattering. Data were recorded at X-ray energies of 9.70 and 7.26 keV. Both energies yielded anomalous difference electron density, indicating that both zinc and iron are found in the active site of the crystallized protein complex, though no metal was added during crystallization (Fig. S3). The E145A and H128A mutants had about 5-7 fold less metal (combined iron and zinc) and 2-3 fold less zinc than the wild-type complex (Fig. 4A, S4B, S4C). However, the D195A complex had as much zinc as the wild-type complex and also contained substantial amounts of iron (Fig. 4A, S4B, S4C). We conclude that D195 is not needed for stable metal binding. Consistent with this observation, in the second complex in the asymmetric unit of the crystal structure, D195 is flipped out towards the solvent, and the metal is instead coordinated by LytH D212 from the neighboring complex, which may be a result of crystal packing (Fig. S1D). More studies will be required to understand the role of D195 in catalysis.

**Fig. 4.**
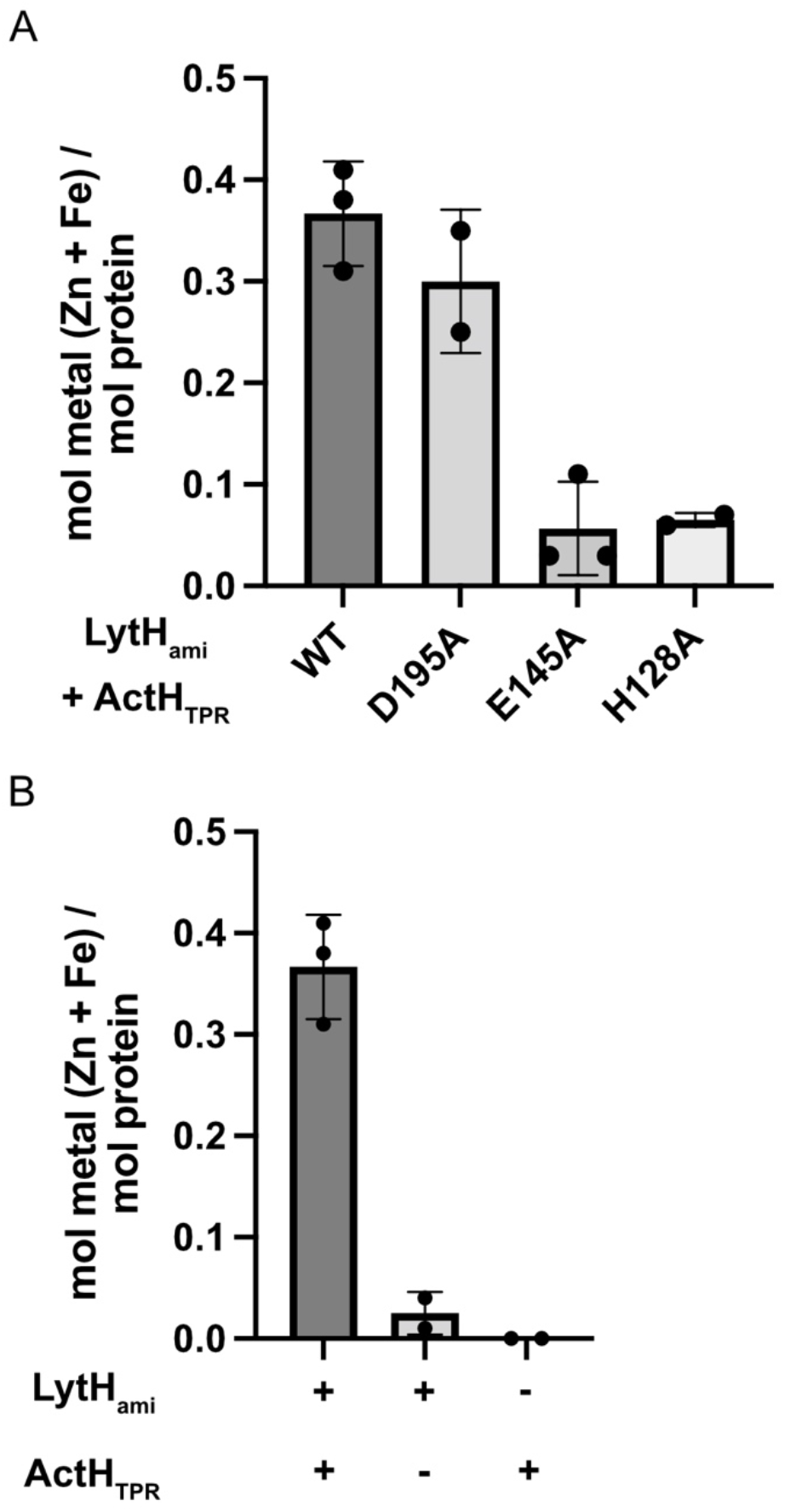
LytH D195 is not required for metal binding, which is stabilized by ActH binding. (*A*) Zinc and iron were quantified in co-purifications of wild-type or mutant LytH_ami_ with ActH_TPR_ by ICP-MS. LytH D195 is not required for metal binding, although H128 and E145 are. Each dot represents an independent purification. (*B*) Zinc and iron were quantified in purifications of the LytH_ami_-ActH_TPR_ complex or the single proteins alone by ICP-MS. Each dot represents an independent purification. Significantly more metal is found in the complex than in LytH_ami_ alone.

### ActH stabilizes metals in the LytH active site

We made an interesting observation while purifying the LytH_ami_-ActH_TPR_ mutant complexes for ICP-MS. When ActH_TPR_ was co-purified from *E. coli* with either wild-type or mutant LytH_ami_ and submitted to size exclusion chromatography, the LytH_ami_ wild-type and D195A complexes eluted as single peaks. However, only small peaks for the complex were seen for LytH_ami_ H128A and E145A, with the majority of the protein eluting as the individual proteins (Fig. S5). This observation suggested that LytH metal binding is necessary for stable complex formation with ActH.

We wondered whether ActH, in turn, stabilizes metal in the active site of LytH. To test this, we measured the amount of zinc and iron in purified samples of the LytH_ami_-ActH_TPR_ complex or individual proteins alone and found that the molar ratio of metal (combined zinc + iron) to protein was about 15-fold higher in the complex than in LytH_ami_ alone (Fig. 4B, S4B, S4C). The LytH_ami_ D195A-ActH_TPR_ complex similarly had significantly more metal than LytH_ami_ D195A alone (Fig. S4A). Only trace amounts of metal were found in purified samples of the ActH TPRs alone. We conclude that ActH stabilizes the binding of metals in the LytH active site, and LytH metal-binding in turn stabilizes the LytH-ActH interface.

### The ActH TPRs are necessary and sufficient for LytH activity in cells

Knowing that the ActH TPRs are sufficient to activate LytH *in vitro*, we next wondered whether they would also suffice for activating LytH in cells. A knockout of *lytH* has a striking phenotype of unusually large cells with division defects due to poorly controlled growth. This mutant is also particularly sensitive to the beta-lactam oxacillin. ActH mutants have similar morphological defects and increased susceptibility to oxacillin (7). We asked whether supplying just the TPRs tethered to the membrane would be sufficient to rescue these cellular defects. In addition to demonstrating sufficiency, such a result would imply that these Δ*actH* phenotypes are due to loss of LytH activity rather than loss of a function of the rhomboid or intracellular domains of ActH. We introduced several FLAG-tagged truncation mutants of ActH on single copy integrative plasmids into a Δ*actH* background and tested growth on oxacillin. A truncation lacking the TPRs was not able to restore growth to the Δ*actH* mutant. However, when the TPRs were fused to a single TM helix of the ActH rhomboid protease domain, whether the first or last helix, growth was comparable to wild-type (Fig. 5A, S6A). We wondered if a single pass TM-TPR construct would also rescue the morphological defects of Δ*actH*. We stained *S. aureus* cells with the membrane dye Nile red and quantified their size. A single pass TM-TPR construct produced cells of wild-type size, whereas the construct lacking the TPRs did not correct the size defect of Δ*actH* cells (Fig. 5B, S7). The chromosomally-integrated FLAG-tagged proteins in all of these strains were undetectable by Western blot. To ensure that the lack of function of the construct lacking the TPRs was not due simply to poor expression, we introduced the same truncations on a plasmid with higher expression levels. All proteins could then be detected by FLAG Western blot (Fig. S6C), but the construct lacking the TPRs still did not restore growth of Δ*actH* on oxacillin (Fig. S6B). Because only the TPRs on a single TM helix are able to restore both normal cell size and resistance to oxacillin, we have concluded that the rhomboid protease and intracellular domains of ActH are dispensable for these phenotypes. Paired with our *in vitro* data, these studies show that the morphological and beta-lactam susceptibility defects of Δ*actH* cells are due to loss of LytH activity and that the ActH TPRs anchored in the membrane are both sufficient and necessary to activate LytH in cells.

**Fig. 5.**
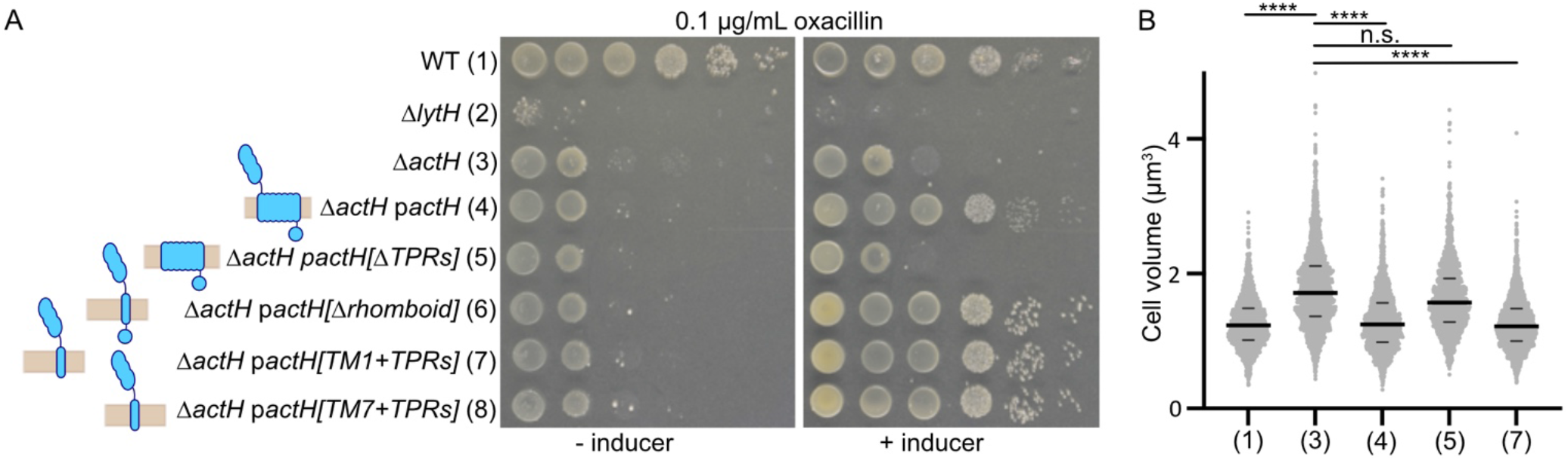
The ActH TPRs on a single transmembrane helix anchor are sufficient for LytH activation in cells. (*A*) *S. aureus* HG003 Δ*lytH* and Δ*actH* are both sensitive to oxacillin. This sensitivity is complemented by ActH truncations with the TPR domain on either the first (row 7) or last (row 8) transmembrane helix of the rhomboid protease domain, but not by a construct lacking the TPR domain (row 5). Strains used are HG003 (1) wild-type, (2) Δ*lytH*, (3) Δ*actH*, (4) Δ*actH pactH*, (5) Δ*actH pactH[2-367]*, (6) Δ*actH pactH[1-178, 365-487]*, (7) Δ*actH pactH[151-178, 365-487]*, and (8) Δ*actH pactH[337-487*]. All constructs were expressed from pTP63 by induction with 0.4 μM anhydrotetracycline. (*B*) *S. aureus* cells were stained with the membrane dye Nile Red, and the volumes of non-dividing cells were estimated and plotted (7). Complementation of Δ*actH* with TPR-containing constructs of ActH correct the size defects associated with loss of LytH activity. The numbers on the x-axis correspond to the strains in subfigure A. In each plot, each dot represents an individual cell, the larger horizontal bars mark the median, and the smaller horizontal bars demarcate the interquartile range. Over 1,500 cells were counted for each sample. The sample medians are (1) 1.234 μm^3^, (3) 1.717 μm^3^, (4) 1.252 μm^3^, (5) 1.573 μm^3^, and (7) 1.220 μm^3^. P-values were determined by two-sided Mann-Whitney U tests. For (1) and (3), p = 1.8 x 10^-204^; for (3) and (4), p = 1.9 x 10^-162^; for (3) and (5), p = 2.0 x 10^-15^; for (3) and (7), p = 1.7 x 10^-298^; **** signifies p < 10^-100^.

### The LytH SH3 domain is required for activity in cells

Like the rhomboid domain of ActH, the SH3 domain of LytH is not required for activity *in vitro*, so we wondered if it would also be dispensable in cells. When we introduced a LytH construct lacking the SH3 domain into a Δ*lytH* background, it was not able to restore growth on oxacillin (Fig. S9C). The SH3 domain is thus essential for LytH activity in cells. DeepMind’s AlphaFold2 (31) predicted the same binding interface between the LytH amidase domain and ActH TPRs seen in our structure and allowed us to visualize the modeled full-length complex (Fig. S8). The extracellular complex sits atop the ActH rhomboid domain, with the SH3 domain of LytH descending on the back of the amidase domain to the single LytH TM helix. We wondered if the SH3 domain is necessary mainly as a spacer to position the catalytic domain for interaction with ActH. To test this, we replaced the native SH3 domain of LytH with either a linker or an SH3 domain from the *Bacillus subtilis* amidase YrvJ (Fig. S9A, S9B), which is also membrane-bound. All constructs were stably expressed (Fig. S9D), but none was able to restore growth on oxacillin (Fig. S9C). Our inability to replace the SH3 domain with a similar domain from another species suggests that it is not simply a spacer. In bacteria, SH3 domains are known to bind and recognize cell wall substrates (32–35). In eukaryotes, where SH3 domains were first identified, they are found in a variety of signaling and cytoskeletal proteins and classically bind to proline-rich peptides (36, 37). Despite its dispensability *in vitro*, the LytH SH3 domain may be important for *in cellulo* substrate recognition, or it may mediate a protein-protein interaction with a yet unidentified binding partner.

## DISCUSSION

This work reveals the mechanism underlying a new class of amidase activators. First, we showed that the extracellular TPR domain of ActH activates LytH both *in vitro* and in cells. The rhomboid and intracellular domains of ActH are dispensable for phenotypes associated with both Δ*lytH* and Δ*actH*, suggesting that ActH has another, unknown function with similar temporal and spatial requirements as LytH activation. Next, we reported a crystal structure of a complex of the extracellular domains of LytH and ActH that shows an unusual mode of TPR binding. We further showed that ActH activates LytH at least in part by stabilizing metals in the LytH active site, a strategy which, as we discuss below, seems to be employed by activators in different structural classes.

Hydrolase activity must be carefully tuned to allow cell growth and division but avoid excessive cleavage of the essential cell wall. Bacteria employ diverse strategies to ensure that hydrolases only act in the correct time and place. These strategies include regulating hydrolase expression, modifying hydrolase substrates in the cell wall, and targeting hydrolases to particular cell wall compartments (5). Direct protein regulators of hydrolases have also been identified (10–14, 38), with the first characterized amidase activators being NlpD and EnvC, which activate the cell separation amidases AmiA/B/C in Gram-negative organisms (10, 39–41). Crystal structures of AmiB and AmiC have shown they contain an alpha helix that blocks the active site and is purportedly displaced upon interaction with the activator (22, 23). LytH does not have a blocking helix. Although we were unable to crystallize the LytH amidase domain alone, the structure of LytH observed in the complex is similar to those of amidases that are functional without an activator. However, we have shown that ActH stabilizes metals in the LytH active site, presumably by inducing small conformational changes.

Recently, the *Clostridioides difficile* lipoprotein GerS was also found to stabilize zinc binding in the amidase CwlD to promote activity (30). Moreover, as with ActH and LytH, metal cofactor binding was found to be important for stable association between GerS and CwlD. GerS bears no resemblance to ActH. Instead, it is a lipoprotein tethered to diacylglycerol in the membrane and has a single extracellular domain largely composed of an anti-parallel beta sheet. That GerS and ActH, two structurally different proteins, both act by stabilizing metal binding in their respective amidases suggests that metal cofactor stabilization is a widespread strategy employed for amidase activation.

Unexpectedly, we found that LytH can bind iron in its active site in place of zinc, and that this binding is also stabilized by ActH. Scattered reports show that other amidases can use metals other than zinc to promote hydrolysis (25, 42). In one study, CwlV from *Paenibacillus polymyxa* was found to purify predominantly with zinc, but also contained significant amounts of manganese. Moreover, CwlV had robust activity when bound to either cobalt or manganese (42). To our knowledge, iron has not been reported as a cofactor in peptidoglycan amidases. However, there are examples of zinc-dependent hydrolases that can use iron(II) in place of zinc(II) to cleave amide bonds. LpxC, which catalyzes the committed step in Lipid A biosynthesis by hydrolyzing an N-acyl bond, can bind either zinc(II) or iron(II) under native conditions, and although the affinity for zinc is greater than for iron, the enzyme is more active with an iron cofactor (43). Because exchangeable intracellular iron(II) is present in greater abundance than zinc(II) under most conditions, it has been argued that iron is the dominant cofactor. Histone deacetylase 8 (HDAC8) was similarly found to use either iron or zinc and has higher activity when bound to iron (44). Although LpxC, HDAC8, and LytH have different substrates, they all cleave amide bonds. It is thus conceivable that LytH similarly makes use of different metal cofactors depending on the conditions.

We have shown that the rhomboid protease domain of ActH is dispensable for the shared phenotypes of Δ*lytH* and Δ*actH*, implying that there is no necessary interaction between the TM helix of LytH and the intramembrane domain of ActH. In contrast, the glucosaminidase SagB, which is also regulated by an intramembrane protein with homology to a family of proteases, forms close contacts through its TM helix with intramembrane helices of the protease, SpdC (38). Protein-protein interactions through membrane domains is a common theme in all cells. That ActH’s hydrolase-regulating function is entirely contained in its extracellular TPR domain, yet ActH homologs with both a rhomboid domain and a TPR domain are widespread in Firmicutes (7, 45), suggests there may be some connection between the LytH-activating role of ActH and its unknown other roles. Rhomboid proteases are found in all domains of life and have important roles in eukaryotes (46), but their functions in bacteria have remained more mysterious (45, 47, 48). Our constructs that lack the rhomboid protease domain, yet activate LytH, now allow exploration of phenotypes specifically associated with the rhomboid domain.

An unanswered question is what purpose is served by having an activator of LytH. A standard view in the field is that cell wall hydrolase activators are required to prevent excessive cleavage of the cell wall. However, some cell wall hydrolases have intrinsic activity, and temporal or spatial mechanisms are used for regulation. For example, the membrane-bound cell wall hydrolase SagB is intrinsically active but is unable to effect peptidoglycan cleavage when SpdC is deleted, evidently because it cannot access substrate that is partially crosslinked into the cell wall matrix unless it is properly presented atop SpdC (38). Investigating the conditions under which ActH and LytH are expressed, and the levels to which they are natively expressed, could be helpful in elucidating the purpose of requiring complexation for activity and might begin to shed some light on why the LytH activation domain is found in a much larger protein with other functions.

## Acknowledgements

We would like to thank Brian Jackson at the Dartmouth Trace Element Analysis Core for help with the elemental analyses. We thank the Microscopy Resources On the North Quad (MicRoN) facility at Harvard Medical School for their training and support. We thank the staff at Advanced Photon Source GM/CA beamlines for support of x-ray data collection. GM/CA@APS is funded by the National Cancer Institute (ACB-12002) and the National Institute of General Medical Sciences (AGM-12006, P30GM138396). The Eiger 16M detector at GM/CA-XSD was funded by NIH grant S10 OD012289. Portions of this research were conducted at the Advanced Photon Source, a U.S. Department of Energy (DOE) Office of Science User Facility operated for the DOE Office of Science by Argonne National Laboratory under Contract No. DE-AC02-06CH11357. SBGrid provided structural biology software support. We would like to acknowledge Bailey Plaman for preliminary work on LytH-ActH binding assays during her rotation, and Liz Nolan and Theodore Betley for helpful discussions of metalloenzymes. Funding for this work was provided by National Institutes of Health grants R01 AI139011 and R01 AI148752 to S.W. and T32 GM007753 and F30 AI156972 to J.E.P., the National Science Foundation DGE1144152 to T.D., and a Merck Postdoctoral Fellowship from the Helen Hay Whitney Foundation to M.A.S.

## Author Contributions

J.E.P., T.D., and S.W. conceived the project. J.E.P., A.C.K., and S.W. designed and coordinated the overall study. J.E.P. performed the biochemical and cellular experiments. J.E.P. and M.A.S. performed crystallographic experiments. T.D. performed preliminary experiments and constructed several strains. The manuscript was written by J.E.P. and S.W. with input from all authors.

## Data availability

Crystallographic data are available from the Protein Data Bank with PDBID 7TJ4.

## Methods

### Materials

Unless otherwise indicated, all chemicals and reagents were purchased from Sigma-Aldrich. Restriction enzymes, KOD DNA polymerase, Q5 2x Master Mix, Phusion 2x Master Mix, and T4 polynucleotide kinase were purchased from New England Biolabs. The In-fusion HD Cloning Plus kit was purchased from Takara Bio USA. Oligonucleotide primers were purchased from Integrated DNA Technologies. Culture media were purchased from Becton Dickinson. *S. pneumoniae* Δ*murMN* Lipid II was isolated from cells as described previously (49, 50). Lipid II was labeled with ATTO488 as previously described (51). *S. aureus* SgtB^Y181D^ was expressed and purified as previously reported (52). Genomic DNA was isolated using a Wizard Genomic DNA Purification kit (Promega).

### Bacterial growth conditions

*E. coli* strains were grown with shaking at 37 °C in lysogeny broth (LB), Terrific Broth (TB), or on agarized LB plates with appropriate antibiotics. *S. aureus* strains were grown with shaking at 30 or 37 °C in tryptic soy broth (TSB) or on agarized TSB plates containing antibiotics as appropriate. Plasmids were cloned using *E. coli* NEB 10-beta cells. *E. coli* Stellar cells were used for cloning with the In-fusion HD Cloning Plus kit. The *E. coli* C43 (DE3) strain was used for overexpression of membrane-anchored proteins, and the BL21 (DE3) strain was used for overexpression of all soluble proteins. The following concentrations of antibiotics were used: carbenicillin, 100 μg/mL; chloramphenicol, 10 μg/mL; erythromycin, 10 μg/mL; kanamycin, 50 μg/mL (neomycin, 50 μg/mL was added as well for kanamycin resistant *S. aureus strains);* tetracycline, 3 μg/mL. The bacterial strains, plasmids and oligonucleotide primers used in this study are summarized in Supplementary Tables. Protocols for plasmid construction can be found in the Supplementary Methods.

### Protein expression

For each soluble protein, *E. coli* BL21(DE3) containing the expression plasmid of interest was grown in 1-1.5 L LB supplemented with the appropriate antibiotics at 37 °C with shaking until OD600 ~0.6. The culture was cooled to 16 °C, and protein expression was induced by adding 500 μM isopropyl β-D-1-thiogalactopyranoside (IPTG). For each membrane-bound protein, *E. coli* C43(DE3) containing the expression plasmid of interest was grown in 1.5 L TB supplemented with appropriate antibiotics at 37 °C with shaking until OD600 ~0.8. The culture was cooled to 16 °C, and protein expression was induced by adding 1 mM IPTG. Cells were harvested 18 h post-induction by centrifugation (4,000 x g, 10 min, 4 °C), and the pellet was stored at −80 °C.

### Purification of soluble Hisθ-tagged proteins

Proteins from expression constructs pTD2 and pTD3 were purified as previously described (7). For elemental analyses, protein expressed from pTD3 was purified as described here. All steps after cell lysis were performed at 4 °C. Cells were resuspended in 30 mL Buffer A (50 mM HEPES pH 7.5, 500 mM NaCl, 10% glycerol) supplemented with 5 mM MgCl2, 1 mM tris(2-carboxyethyl)phosphine (TCEP), 1 mg/mL lysozyme, 250 μg/mL DNase, 1 mM phenylmethylsulfonyl fluoride (PMSF) and Roche cOmplete Protease Inhibitor and stirred to homogenize. The resuspended cells were then passaged through a cell disruptor (EmulsiFlex-C5, Avestin) at 15,000 psi three times to lyse. Cell debris was removed by centrifugation (10,000 x g, 5 min, 4 °C), and the membrane fraction was removed by ultracentrifugation of the supernatant (119,000 x g, 45 min, 4 °C). The resulting supernatant was supplemented with 1 mL pre-equilibrated Ni-NTA resin (Qiagen) and 10 mM imidazole and stirred for 30 min at 4 °C. The sample was then loaded onto a gravity column and washed with 30 mL Buffer A containing 10 mM imidazole, 30 mL Buffer A containing 20 mM imidazole, and 30 mL Buffer A containing 40 mM imidazole. The protein was then eluted in 20 mL Buffer A containing 300 mM imidazole. The eluate was further purified by size exclusion chromatography (SEC) with a Superdex 75 10/300 GL (for expression constructs pJP62, pJP151, and pJP152) or Superdex 200 Increase 10/300 GL (all others, pJP62 for elemental analysis) equilibrated in Buffer A. Fractions containing the target protein were concentrated by centrifugal filtration. The absorbance at 280 nm was measured using a NanoDrop One Microvolume UV-Vis Spectrophotometer (ThermoFisher Scientific), and the predicted extinction coefficient was used to calculate concentration. Protein samples were then aliquoted and stored at −80 °C.

### Purification of membrane bound His-tagged proteins

Full length His6-ActH (construct pTD52) was purified as previously described (7). Full length LytH-His6 (construct pTD42) was purified as described for His-tagged soluble proteins with the following modifications. After ultracentrifugation, the membrane fraction was collected and resuspended in 30 mL solubilization buffer (Buffer A + 1% (w/v) n-dodecyl β-D-maltoside (DDM) and 1 mM TCEP). The resulting mixture was stirred for 1 hr at 4 °C before ultracentrifugation (119,000 x g, 35 min, 4 °C). The resulting supernatant was supplemented with 0.75 mL pre-equilibrated TALON resin (Takara Clontech) and 1 mM imidazole and stirred for 30 min at 4 °C. The sample was then loaded onto a gravity column and washed with 20 mL each of Buffer A supplemented with 2 mM imidazole/1% DDM, 4 mM imidazole/0.2% DDM, 6 mM imidazole/0.1% DDM, 8 mM imidazole/0.05% DDM, 10 mM imidazole/0.05% DDM, and 15 mM imidazole/0.05% DDM. The protein was then eluted in 10 mL Buffer A containing 0.05% DDM and 150 mM imidazole. The eluate was further purified by size exclusion chromatography (SEC) with a Superdex 200 10/300 GL column equilibrated in Buffer A with 0.05% DDM.

### Purification of soluble FLAG-tagged proteins

All steps after cell lysis were performed at 4 °C. Cells were resuspended in 30 mL Buffer B (50 mM HEPES pH 7.5, 150 mM NaCl, 10% glycerol) supplemented with 5 mM MgCl2, 1 mg/mL lysozyme, 250 μg/mL DNase, and 1 mM phenylmethylsulfonyl fluoride (PMSF) and stirred to homogenize. The resuspended cells were then passaged through a cell disruptor (EmulsiFlex-C5, Avestin) at 15,000 psi three times to lyse. Cell debris was removed by centrifugation (10,000 x g, 5 min, 4 °C), and the membrane fraction was removed by ultracentrifugation of the supernatant (119,000 x g, 45 min, 4 °C). The resulting supernatant was then loaded onto a gravity column with 1 mL α-FLAG G1 affinity resin (Genscript), and the flow through was passed through the column four more times. The resin was washed with 3 x 15 mL of Buffer B, and the protein was eluted with 10 mL of Buffer A supplemented with 0.2 mg/mL FLAG peptide (Genscript). The eluate was further purified by size exclusion chromatography (SEC) with a Superdex 200 Increase 10/300 GL equilibrated in Buffer A. Fractions containing the target protein were concentrated by centrifugal filtration. The absorbance at 280 nm was measured using a NanoDrop One Microvolume UV-Vis Spectrophotometer (ThermoFisher Scientific), and the predicted extinction coefficient was used to calculate concentration. Protein samples were then aliquoted and stored at −80 °C.

### In-gel detection of amidase activity

ATTO488-labeled Lipid II (1.4 μM) was polymerized with 1.8 μM SgtB^Y181D^, a monofunctional peptidoglycan glycosyltransferase with impaired processivity (52), in 1.1x reaction buffer (1x buffer = 50 mM Hepes pH 7.5, 10 mM CaCl_2_, 60 μM Zn(OAc)_2_, and 15% DMSO) at room temperature for 2 h. The polymerization reaction was heat-quenched at 95 °C for 5 min. After cooling, the digestion reaction was set up by adding 1 μL of 5 μM enzyme to 9 μL of the polymerization reaction product (total volume 10 μL). For reactions testing pairs of proteins (LytH + ActH), mixes containing 5 μM of each protein were first prepared and incubated on ice for 20 min before addition to the polymerization reaction product. After incubating the reaction mixtures at room temperature for 5 hours, the reactions were quenched by adding 10 μL 2x Laemmli sample buffer (Bio-Rad). The samples were then loaded onto a 4-20% Mini-PROTEAN TGX Precast Protein gel (Bio-Rad) and run at 180 V. The gels were imaged using a Typhoon FLA 7000 imager.

### Crystallization and structure determination

pJP85 was transformed into BL21 cells, and LytH[117-291]-ActH[365-479] was expressed and purified as described for His-tagged soluble proteins except that the final protein was exchanged into 50 mM Hepes pH 7.5, 150 mM NaCl on the Superdex200 Increase 10/300 GL column. Final protein was aliquoted and flash frozen. Crystals were obtained in a 1:1 ratio of 12.6 mg/mL protein solution to 0.17 M sodium acetate, 0.085 M Tris: HCl pH 8.5, 25.5% (w/v) PEG4000, 15% (v/v) glycerol after 1-2 days at 20 °C. Crystals were harvested with nylon loops and then flash cooled in liquid nitrogen.

Diffraction data were collected at 1.033 Å and 100 K at the Advanced Photon Source GM/CA beamline 23ID-B. Data were collected at 1° / second with ten-fold attenuation, and 0.2° oscillation range. Efforts to solve the structure from this data revealed overlapping lattice patterns from twinned crystals, and the data could not be deconvoluted. Preliminary molecular replacement solutions using Phaser (53) through the Phenix Software Suite (54) with an amidase from *C. difficile* (PDB ID: 4RN7) as a search model demonstrated a trimer forming between three units of LytH, with interactions between helices spanning residues 179-185 and 242-247 on one subunit and a loop from residues 196-203 on the other. In order to disrupt this interaction and force the protein to crystallize in a different lattice, we prepared a series of constructs with mutations in these regions, ultimately solving the structure of LytH[117-291, R245A]-ActH[365-479].

pJP107 was transformed into BL21 cells, and LytH[117-291, R245A]-ActH[365-479] was expressed and purified as described for His-tagged proteins except that the final protein was exchanged into 50 mM Hepes pH 7.5, 150 mM NaCl on the Superdex200 Increase 10/300 GL column. Final protein was aliquoted and flash frozen. Crystals were obtained in a 2:1 ratio of 12.6 mg/mL protein solution to 0.1 M NH4NO3 pH 6.3, 22% (w/v) PEG3350 after 3-7 days at 20 °C. Crystals were harvested with nylon loops after cryoprotection in 0.1 M NH4NO3 pH 6.3, 22% (w/v) PEG3350, 15% glycerol and then flash cooled in liquid nitrogen.

Diffraction data were collected at 1.033 Å and 100 K at the Advanced Photon Source GM/CA beamline 23ID-D. Data were collected at 1° / second with ten-fold attenuation, and 0.2° oscillation range. Data were processed with XDS (55). A complete dataset was obtained from one crystal and processed in space group P222. The structure was solved by molecular replacement using Phaser (53) through the Phenix Software Suite (54) using a *C. difficile* amidase (PDB ID: 4RN7) as a search model (56). A model of LytH and ActH was built with Phenix AutoBuild (57). Iterative rounds of model building and refinement were carried out using Coot (58) and Phenix.refine (59) with automated translation/liberation/screw group selection. Structures were validated with MolProbity (60). Figures were prepared using Pymol (Schrodinger, LLC. The PyMOL Molecular Graphics System. Version 2.3.4). All structural biology software was accessed through SBGrid (61). The protein interface was analyzed using the ‘Protein interfaces, surfaces and assemblies’ service PISA at the European Bioinformatics Institute (http://www.ebi.ac.uk/pdbe/prot_int/pistart.html).

To examine the identity of the metal in the active site of LytH we collected Friedel-pair data at X-ray energies of 9.70 and 7.26 keV. Data for isomorphous crystals was processed with XDS and phased through rigid-body refinement in Refmac5 (62).

### *In vitro* pull-down binding assay

Pairs of FLAG-tagged LytH proteins and His-tagged ActH proteins were mixed 1:1 at a final concentration of 11 μM for each and incubated on ice for 10 min (P, pre-loading). This mix (16 μL) was then loaded onto 15 μL of α-FLAG G1 resin (Genscript) pre-equilibrated in FLAG resin buffer (50 mM Hepes pH 7.5, 150 mM NaCl, 10% glycerol) in a microspin column (Pierce), and the flow-through was collected. The resin was then washed three times with 2 CVs of FLAG resin buffer each time, incubating on ice for 5 min with each wash before collecting. The protein was then eluted with 1 CV of FLAG resin buffer supplemented with 0.2 mg/mL FLAG peptide (Genscript) after incubating with the elution buffer for 5 min on ice. A 5x Laemmli buffer was added to each sample, and the samples were then loaded onto a 4-20% Midi-PROTEAN TGX Precast Protein gel (Bio-Rad) and run at 180V. The gel was stained with Instant Blue (abcam) and imaged.

### Elemental analyses

SEC-purified proteins at 20 μM in 50 mM Hepes pH 7.5, 500 mM NaCl, 10% glycerol were diluted 7X in water and analyzed by ICP-MS (8900, Agilent, Santa Clara, CA) in helium mode at the Dartmouth Trace Element Analysis Core.

### *S. aureus* strain construction

To construct strains containing pTP63 plasmids, the plasmids were first electroporated into TD011, and transformants were selected on TSA supplemented with 10 μg/mL chloramphenicol. pTP63 constructs were then transduced into strain TD177 to produce strains JP299, JP331, JP332, JP334, and JP335.

To construct strains containing ActH or its truncations on pLOW plasmids, the plasmids were first electroporated into RN4220 wild-type, and the transformants were selected on TSA supplemented with 10 μg/mL erythromycin. pLOW constructs were then transduced into strain TD177 to produce strains JP367, JP368, JP373, JP374, and JP375.

To construct strains containing pLOW plasmids in a Δ*lytH::kan^R^* background, the plasmids were isolated from *E. coli* DC10B and then directly electroporated into TD024 to produce strains TD156, JP391, JP392, JP400, and JP401.

### Spot dilution assays

*S. aureus* cultures in TSB with antibiotics as appropriate were grown overnight at 30 °C with aeration. Overnight cultures were diluted 1:100 into fresh TSB without antibiotics and grown to mid-log phase. The cultures were normalized, five 10-fold dilutions were prepared in TSB, and 5 μL of each dilution were spotted onto TSA plates with or without 0.1 μg/mL oxacillin and inducer. Plates were incubated overnight at 37 °C. A Nikon D3400 DSLR camera fitted with an AF-S Micro-Nikkor 40 mm 1:2.8G lens was used to take pictures of the plates.

### α-FLAG Western blots

*S. aureus* strains were inoculated in TSB with antibiotics as appropriate and the cultures were grown at 30 °C overnight with aeration. Overnight cultures were diluted 1:100 into fresh TSB with or without 1 mM IPTG and grown for 3.5 hr with aeration at 30 °C. For strains containing pLOW constructs, TSB was supplemented with erythromycin. The cultures were normalized, harvested, and lysed in 1x PBS pH 7.4 supplemented with 100 μg/mL lysostaphin, 20 μg/mL DNase, and 5 mM MgCl2 with incubation at 37 °C for 1 hr. Laemmli buffer was then added, and the samples were incubated at 37 °C for an additional 30 min. The samples were then loaded onto a 4-20% PROTEAN TGX Precast Protein gel (Bio-Rad) and run at 180V, transferred to a nitrocellulose membrane (Bio-Rad), and blocked in 1x TBST containing 5% Blotting Grade Blocker (Bio-Rad) for 1 hr at room temperature. Membranes were then blotted with 1:2000 α-FLAG M2-HRP (Sigma Aldrich A8592) in TBST with 5% Blotting Grade Blocker for 1 hr at room temperature, washed with TBST, and exposed with ECL reagent (Pierce).

### Microscopy analysis of *S. aureus* cells

*S. aureus* cultures were grown overnight at 30 °C in TSB with antibiotics as appropriate. The overnight cultures were then diluted to a starting OD_600_ of 0.02 in 3 mL TSB with 0.4 μM anhydrotetracycline, grown at 37 °C with aeration to mid-log phase, and normalized. Cells (1 mL normalized culture) were then labeled with 5 μg/mL Nile red for 5 min at 37 °C with shaking (500 rpm). The cells were pelleted (4,000xg, 2 min), most of the supernatant was removed, and the cells were resuspended in the remaining supernatant (about 50 μL). Cells were spotted onto a thin 2% agarose pad prepared in 1x PBS pH 7.4 and covered with a No. 1.5 cover slip, and sealed with Valap (equal weights of petroleum jelly, lanolin, and paraffin). Brightfield, phase-contrast, and widefield epifluorescence microscopy images were obtained using a Nikon Ti inverted microscope fitted with a custom-made cage incubator set at 30 °C, a Nikon motorized stage with an OkoLab gas incubator and a slide insert attachment, either an Andor Zyla 4.2 Plus sCMOS or a Hamamatsu Orca Flash 4.0 V3 camera, Lumencore SpectraX LED Illumination, Plan Apo lamba x100/1.45 Oil Ph3 DM objective lens, and Nikon Elements 4.30 acquisition software. The microscope was fitted with a 49008 Chroma ET filter cube for detecting Nile red. Exposure times for Nile red labeling were 20-80 ms. Images were analyzed using FIJI (63) and Matlab scripts developed in-house. *S. aureus* cell volumes were estimated using StaphSizer as previously described (7). Only cells without a visible septum were included for this analysis.

