## Supplementary Information for "Metal cofactor stabilization by a partner protein is a widespread strategy employed for amidase activation"

| | LytH R245A ActH | LytH R245A ActH | LytH R245A ActH |
|---|---|---|---|
| **Data Collection** | | | |
| Wavelength (Å) | 1.033 | 1.278 | 1.707 |
| Space group | $P22_12_1$ | $P22_12_1$ | $P22_12_1$ |
| *a*, *b*, *c* (Å) | 44.6, 71.3, 190.4 | 44.7, 71.2, 190.9 | 44.7, 71.5, 190.9 |
| α, β, γ (°) | 90, 90, 90 | 90, 90, 90 | 90, 90, 90 |
| Resolution (Å) | 43.4 - 1.8 (1.86 – 1.8) * | 47.5-2.1 (2.14-2.1) | 47.5-2.5 (2.56-2.5) |
| $R_{merge}$ (%) | 14.2 (381.2) | 21.3 (295.2) | 13.4 (85.4) |
| $CC_{1/2}$ (%) | 99.8 (45.0) | 99.1 (65.2) | 99.4 (63.4) |
| $I / \sigma I$ | 12.0 (0.82) | 8.5 (1.3) | 10.5 (2.0) |
| Completeness (%) | 97.0 (93.6) | 97.4 (91.2) | 96.9 (91.1) |
| Multiplicity | 13.1 (12.8) | 6.7 (6.9) | 6.9 (6.5) |
| **Refinement** | | | |
| Resolution (Å) | 43.4 – 1.8 | | |
| No. reflections | 55,647 | | |
| $R_{work}$ / $R_{free}$ (%) | 18.2/22.3 | | |
| No. atoms | 5039 | | |
| Protein | 4717 | | |
| Ligand | 2 | | |
| Water | 320 | | |
| *B*-factors | 42.6 | | |
| Protein | 42.7 | | |
| Ligand | 32.7 | | |
| Water | 41.8 | | |
| R.m.s. deviations | | | |
| Bond lengths (Å) | 0.010 | | |
| Bond angles (°) | 0.95 | | |
| Ramachandran | | | |
| Favored (%) | 97.3 | | |
| Allowed (%) | 2.7 | | |
| Outlier (%) | 0 | | |

*Values in parentheses are for highest-resolution

**Supplementary Table 1. Crystallographic data collection and refinement statistics for LytH[117-291, R245A]-ActH[365-479].**

A

```
    E. coli AmiC 185 DRPIVIMLDPGHGGEDSGAVGK--YKTREKDVVLQIARRLRSLIEKEGNMKVYMTRNEDI 242
  S. aureus LytH 117 LQGKTIVLDPGHGGSDQGASSNTKYKSLEKDYTLKTAKELQRTLE-KEGATVKMTRTDDT 175
P. polymyxa CwlV 319 NGKKVVVIDAGHGAKDSGAVGIS-RKNYEKTFNLAMALKVESILKQNPKLEVVLTRSDDT 377
                         .:::* ***..*.** .    *. **   *  *  .:.  :: :    * :**.:*

    E. coli AmiC 243 FIPLQVRVAKAQKQRADLFVSIHADAFTSRQPSGSSVFALSTKGATSTAAKYLAQTQNAS 302
  S. aureus LytH 176 YVSLENR-----DIKGDAYLSIHNDALESSNANGMTVYWYHDNQRA-------------- 216
P. polymyxa CwlV 378 FLELKQRVKVAENLKANVFVSIHANSSGSSASNGTETYYQRSASKA-------------- 423
                     :: *: *     . :.: ::*** ::  *   .*  .:       :

    E. coli AmiC 303 DLIGGVSKSGDRYVDHTMFDMVQSLTIADSLKFGKAVLNKLGKINKLHKNQVEQAGFAVL 362
  S. aureus LytH 217 --------------------------------LADTLDATIQKKGLLSNRGSRQENYQVL 244
P. polymyxa CwlV 424 --------------------------------FANVMHKYFAPATGLTDRGIRYGNFHVI 451
                                                     :...:   :     * ..  .  .: *:

    E. coli AmiC 363 KAPDIPSILVETAFISNVEEERKLKTATFQQEVAESILAGIKAYFADGATLARRG      417
  S. aureus LytH 245 RQTKVPAVLLELGYISNPTDETMIKDQLHRQILEQAIVDGLKIYFSA--------      291
P. polymyxa CwlV 452 RETTMPAVLLEVGYLSNAKEEATLFDEDFQNRVAQGIADGITEYLDVK-------      499
                     :   :*::*:* .::**  :*  :    .:: : :.*  *:. *:
```

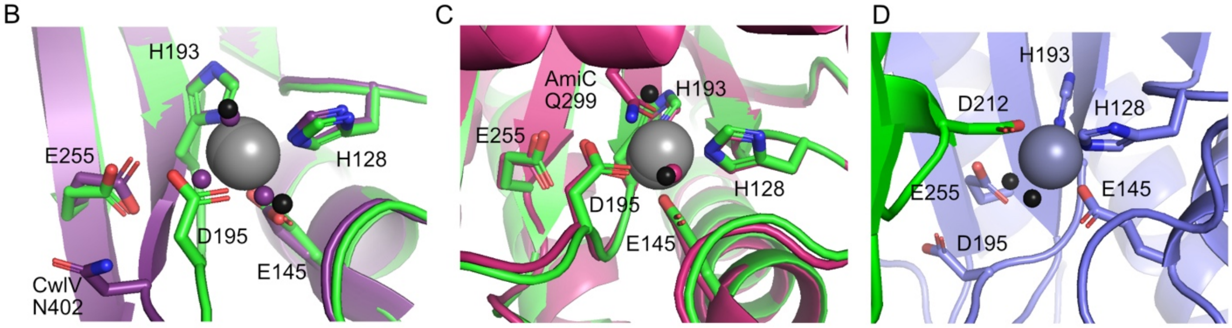


**Supplementary Figure 1. Four amino acid side chains and two waters coordinate a metal ion in the LytH active site.** (*A*) Sequence alignment of the LytH amidase domain with two other amidase_3 domain proteins, *Escherichia coli* AmiC and *Paenibacillus polymyxa* CwlV. Alignments were made using ClustalO (https://www.ebi.ac.uk/Tools/msa/clustalo/). Residues displayed in panels B-D are highlighted in yellow. (*B*) In *Paenibacillus polymyxa* CwlV (purple, PDB 1JWQ) and many other amidase_3 family members, the residue equivalent to LytH (green) D195 is instead an asparagine oriented out towards the solvent. The black balls represent water in the LytH structure, and the purple balls indicate water in the CwlV structure. (*C*) Like LytH (green), AmiC (pink, PDB 4BIN) contains aspartate at the equivalent position to D195, and this aspartate coordinates zinc. The black balls represent water in the structure of LytH and the pink ball water in AmiC. (*D*) In a second complex (blue) in an asymmetric unit in our structure, LytH D195 is flipped out towards the solvent, and zinc is instead coordinated by LytH D212 from the neighboring unit (green).

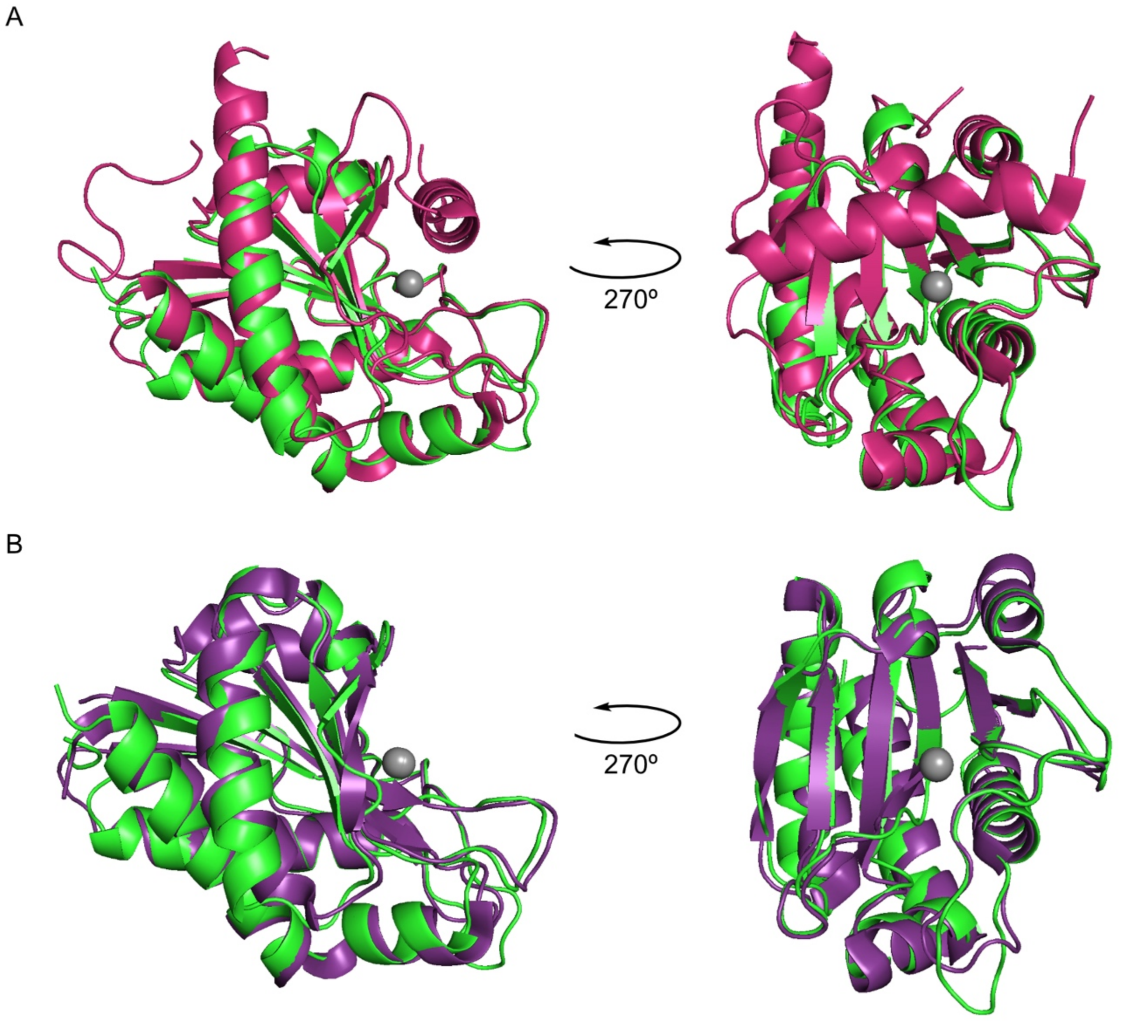


**Supplementary Figure 2. The LytH amidase domain fold is highly conserved with other amidase_3 family proteins.** (*A*) Alignment of the LytH amidase domain (green) with the amidase domain of *E. coli* AmiC (pink, PDB 4BIN). AmiC has an additional helix that blocks its active site, rendering it inactive on its own. (*B*) Alignment of the LytH amidase domain (green) with that of *P. polymyxa* CwlV (purple, PDB 1JWQ), which does not require an activator.

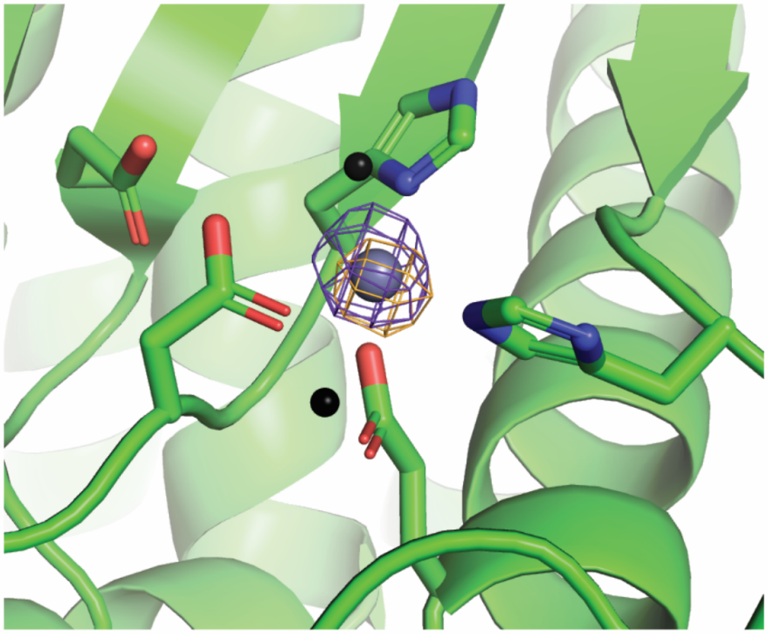

**Supplementary Figure 3. Anomalous scattering at metal center.** Anomalous difference density (5σ) collected above the Zinc K edge 1.278 Å (purple mesh) and Iron K edge 1.707 Å (orange mesh). The peak heights of the two metal sites in the asymmetric unit at 1.278 Å are 7.8 and 7.8 σ, whereas the peak heights of the two metal sites at 1.707 Å are 7.2 and 6.3 σ.

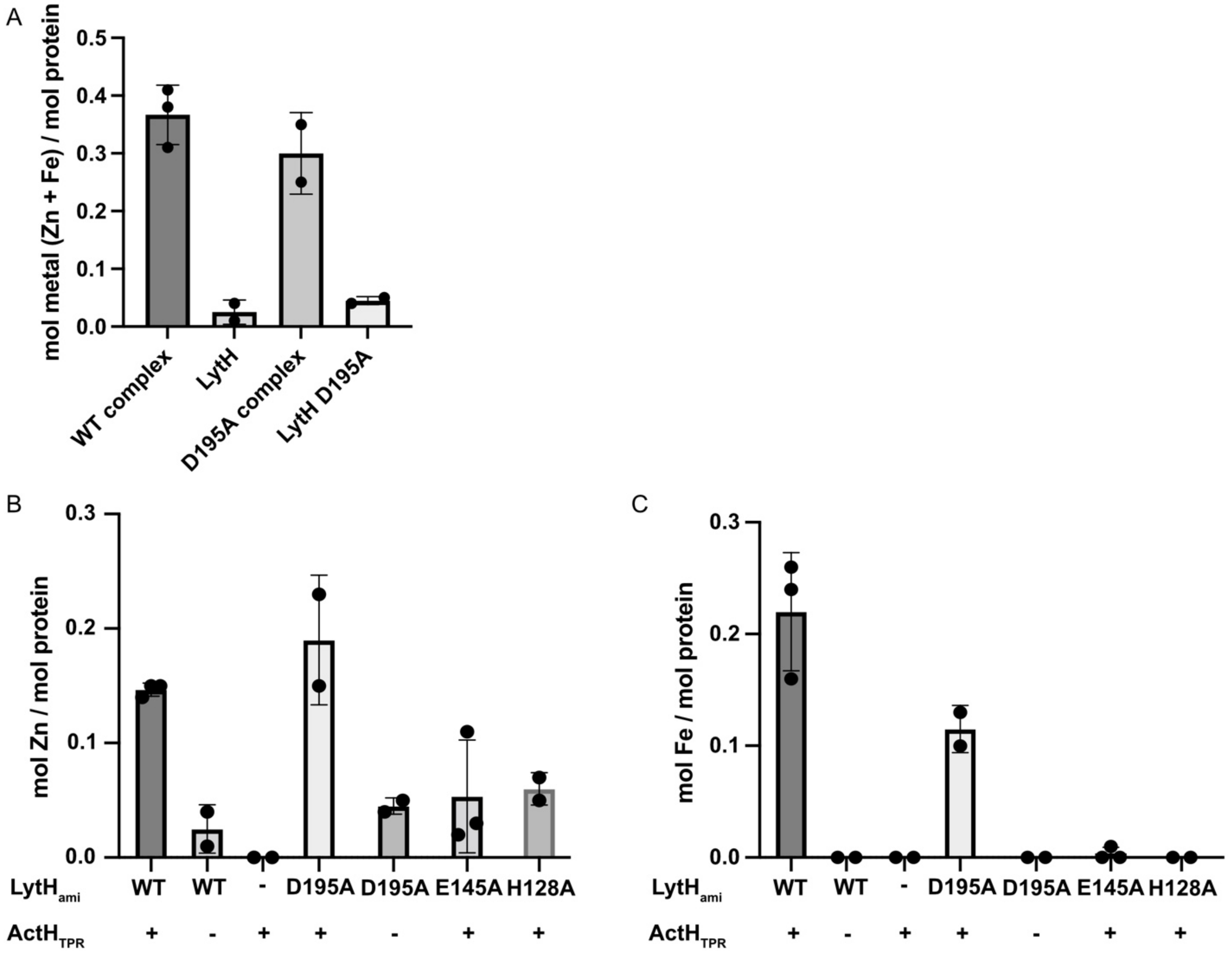


**Supplementary Figure 4. ActH stabilizes metals in LytH.** (*A*) Zinc and iron were quantified in purifications of $LytH_{ami}$-$ActH_{TPR}$ complexes or $LytH_{ami}$ alone by ICP-MS. Each dot represents an independent purification. Significantly more metal is found in the complex than in $LytH_{ami}$ alone, and the same pattern is seen for $LytH_{ami}$ D195A. (*B*) Quantities of zinc measured by ICP-MS are shown for all samples. (*C*) Quantities of iron measured by ICP-MS are shown for all samples.

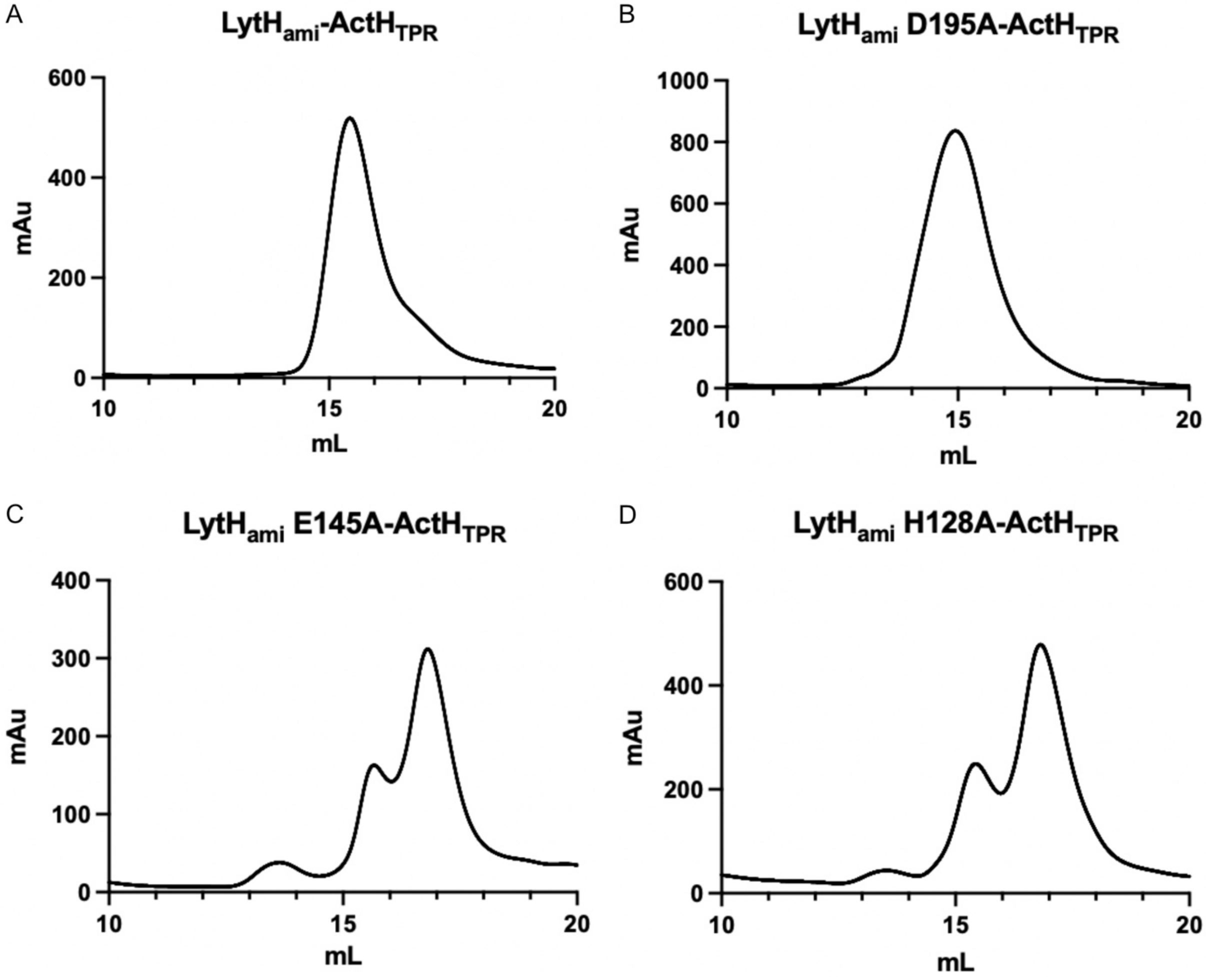


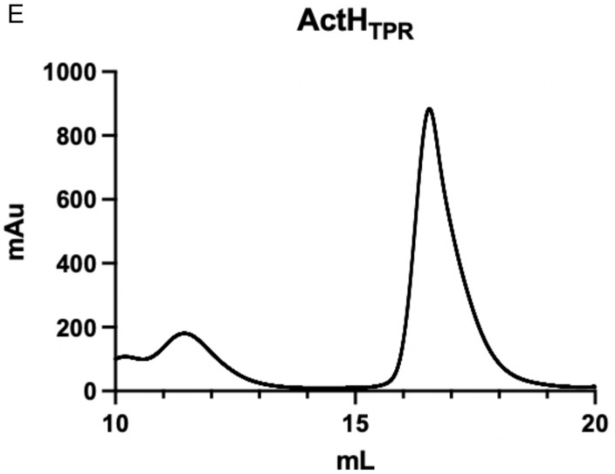


**Supplementary Figure 5. LytH mutants defective in metal binding do not form stable complexes with ActH.** Wild-type or mutant FLAG-tagged LytH$_{ami}$ and His-tagged ActH$_{TPR}$ were co-expressed in *E. coli*, co-purified by nickel-NTA affinity chromatography, and subjected to size exclusion chromatography. Size exclusion chromatograms for co-purifications of (*A*) LytH$_{ami}$-ActH$_{TPR}$, (*B*) LytH$_{ami}$ D195A-ActH$_{TPR}$, (*C*) LytH$_{ami}$ E145A-ActH$_{TPR}$, and (*D*) LytH$_{ami}$ H128A-ActH$_{TPR}$ are shown. While the wild-type and LytH D195A complexes elute as single peaks, the samples with LytH E145A and LytH H128A that have compromised metal binding elute over multiple peaks representing free ActH (~17 mL; see panel (*E*)) and other species.

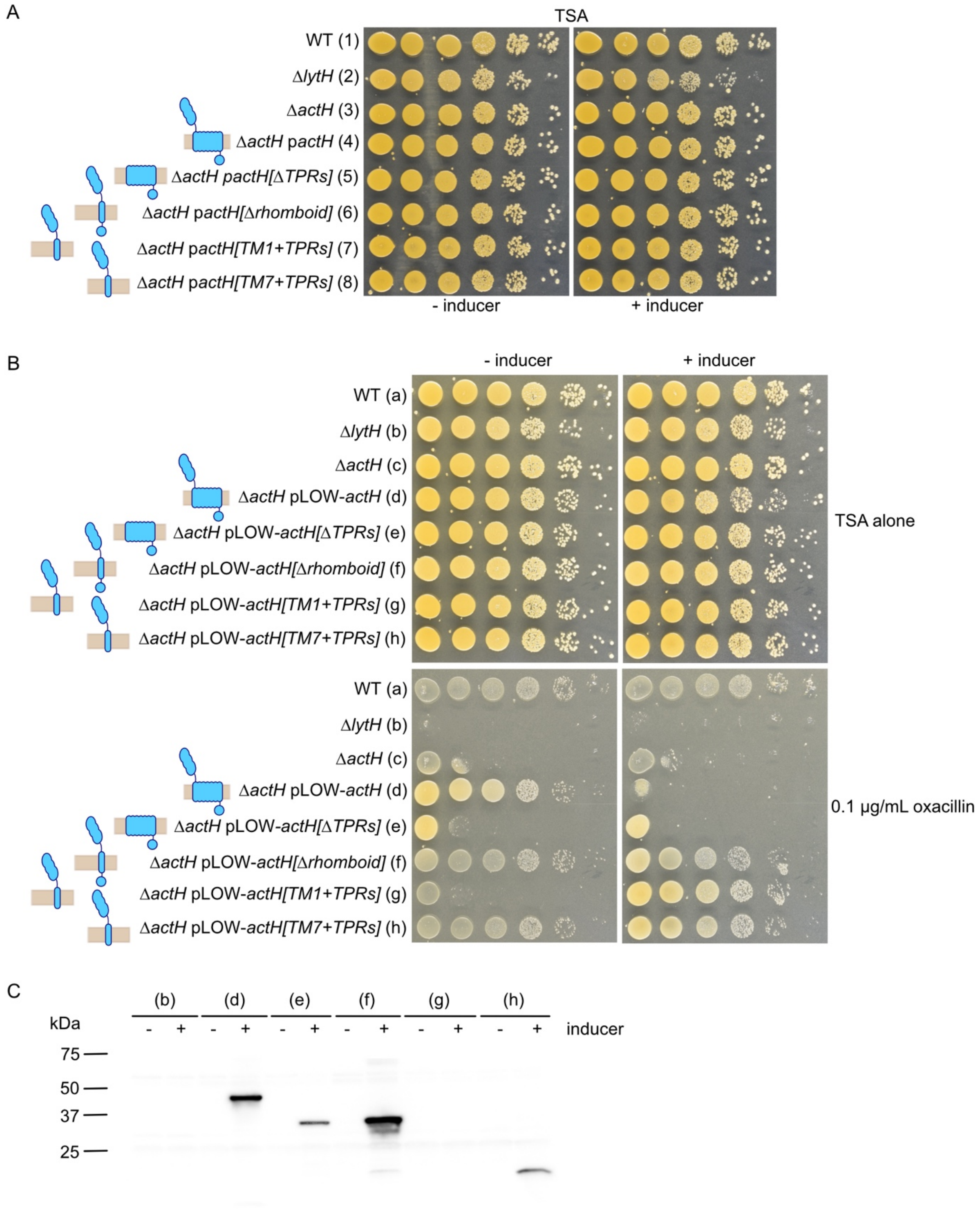


**Supplementary Figure 6. The TPRs of ActH are required for LytH activation in cells.** (*A*) Spot dilutions of strains from Fig. 5A without oxacillin. Strains used are HG003 (1) wild-type, (2) Δ*lytH*, (3) Δ*actH*, (4) JP299 Δ*actH* pTP63-actH-FLAG, (5) JP331 Δ*actH* pTP63-FLAG-actH[2-367], (6) JP332 Δ*actH* pTP63-FLAG-actH[1-178, 365-487], (7) JP334 Δ*actH* pTP63-actH[151-178, 365-487]-FLAG, and (8) JP335 Δ*actH* pTP63-actH[337-487]-FLAG. All ActH constructs were induced with 0.4 µM anhydrotetracycline. (*B*) Spotting assays of strains expressing ActH truncations on pLOW plasmids in an

HG003 Δ*actH* background induced with 1 mM IPTG. Strains used are HG003 (a) wild-type, (b) Δ*lytH*, (c) Δ*actH*, (d) JP373 Δ*actH* pLOW-actH-FLAG, (e) JP367 Δ*actH* pLOW-FLAG-actH[2-367], (f) JP368 Δ*actH* pLOW-actH[1-178, 365-487]-FLAG, (g) JP374 Δ*actH* pLOW-actH[151-178, 365-487]-FLAG, and (h) JP375 Δ*actH* pLOW-actH[337-487]-FLAG. Notably, there is leaky expression from the pLOW vector in the absence of inducer, allowing complementation for some constructs even without inducer. The construct lacking the TPRs still fails to restore growth of Δ*actH* on oxacillin at these higher expression levels, consistent with the TPRs being required for ActH to activate LytH in cells. Interestingly, we noticed that when the full-length ActH protein was expressed at these higher levels, it became toxic in the Δ*actH* background on oxacillin. The construct lacking the rhomboid protease domain but retaining the cytoplasmic domain and the TPRs with a single transmembrane helix (f) did not show toxicity, despite equivalent or higher expression levels (see panel C), suggesting that this phenotype may be due to the rhomboid protease domain. (*C*) α-FLAG Western blot of the strains from (*B*). The construct lacking the TPRs (e) is expressed to equivalent or higher levels than complementing constructs.

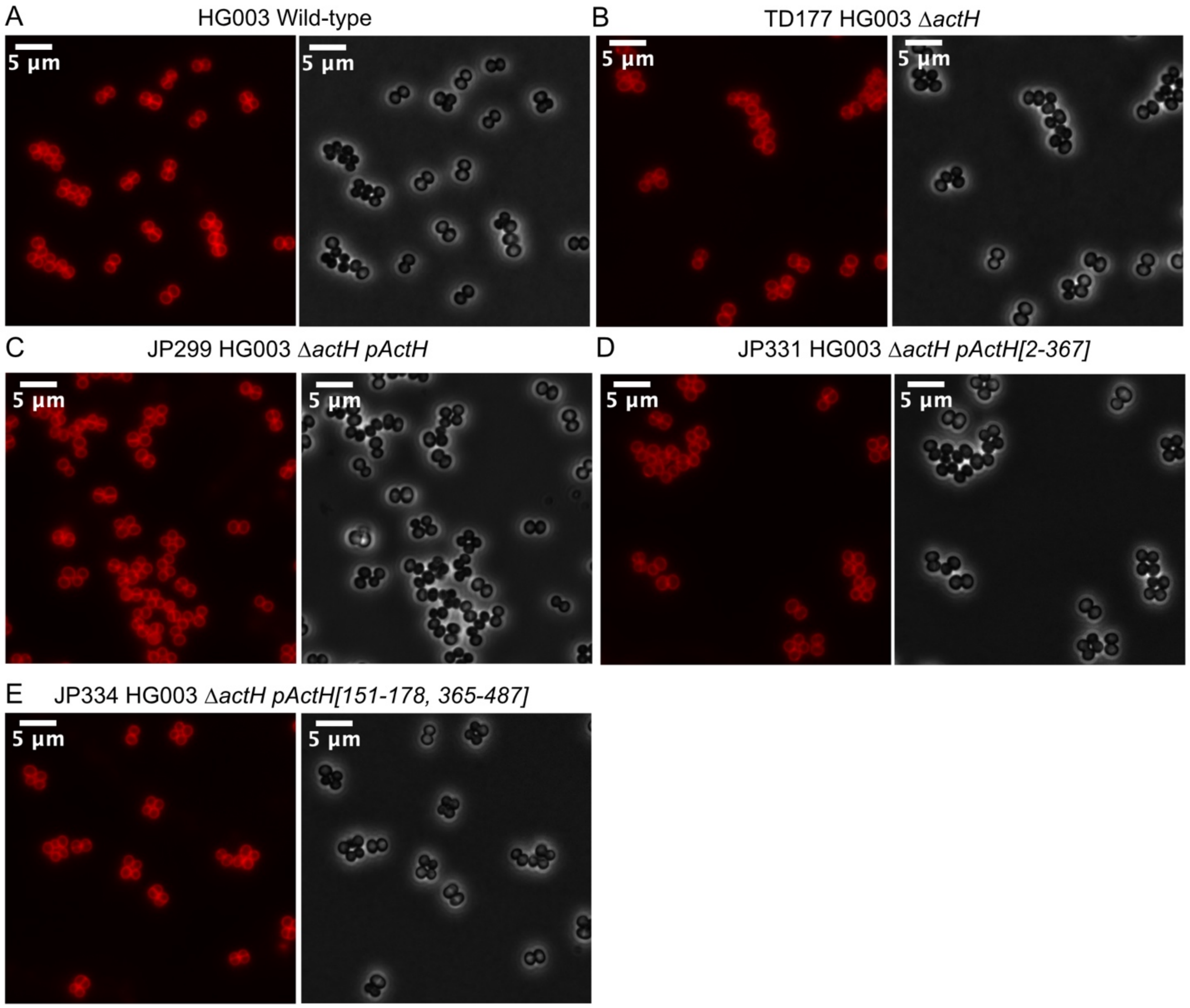


**Supplementary Figure 7. TPR-containing constructs correct the size defects of Δ*actH* cells.** Representative images of *S. aureus* HG003 cells stained with the membrane dye Nile red. (*A*) Wild-type (*B*) TD177 Δ*actH* (*C*) JP299 Δ*actH* pActH (*D*) JP331 Δ*actH* pActH[2-367] (*E*) JP334 ΔactH pActH[151-178, 365-487].

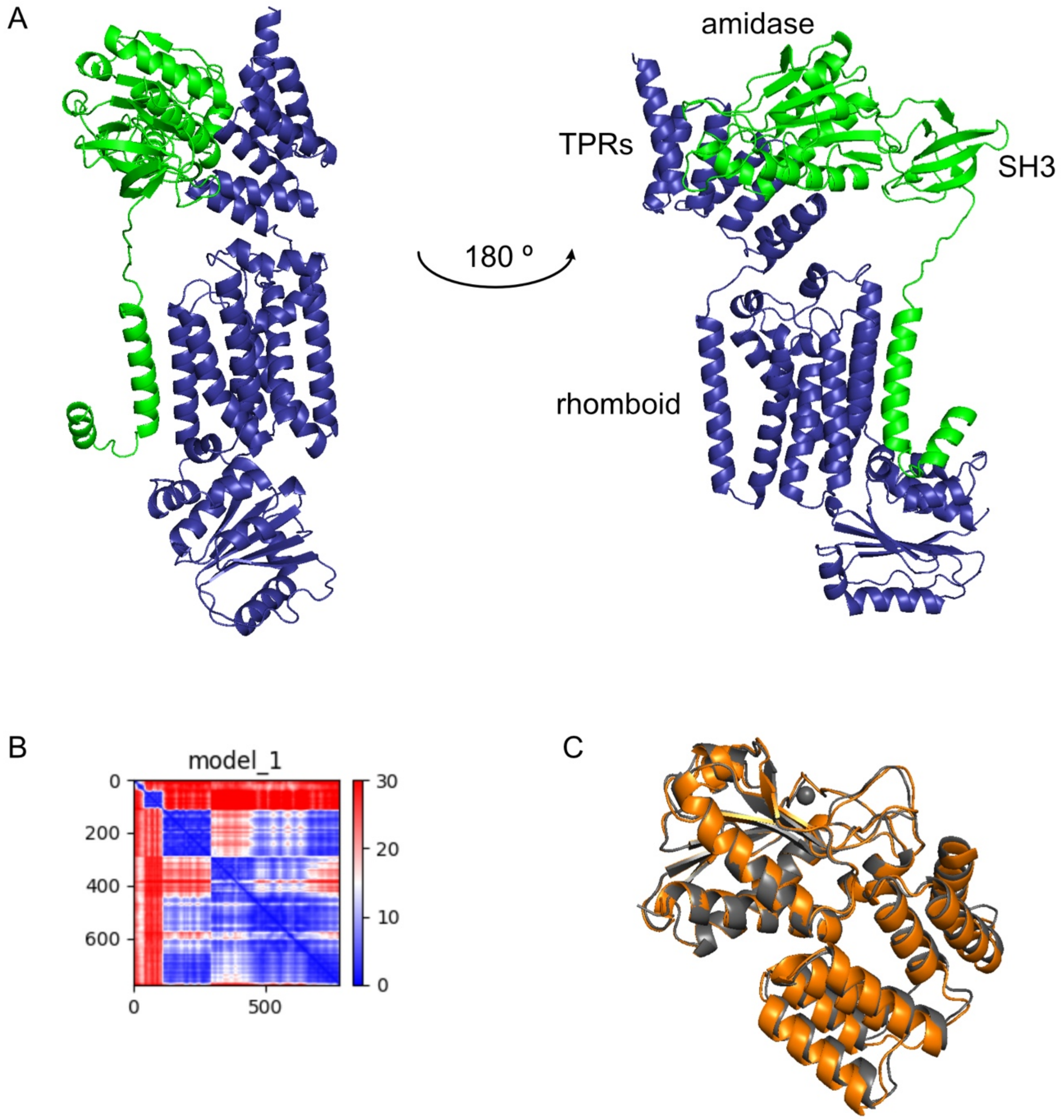


**Supplementary Figure 8. AlphaFold2 predicts binding between the LytH amidase domain and ActH TPRs as seen in our structure and allows modeling of the full protein complex.** (*A*) AlphaFold2 prediction for the complex between the full-length LytH (green) and ActH (blue) proteins. (*B*) Predicted aligned error for this AlphaFold2 model, in which LytH makes up the first 291 residues and ActH the subsequent 487. (*C*) Alignment of the AlphaFold2 predicted structure (orange) and our crystal structure (gray), with RMSD = 0.423 Å over 1120 atoms.

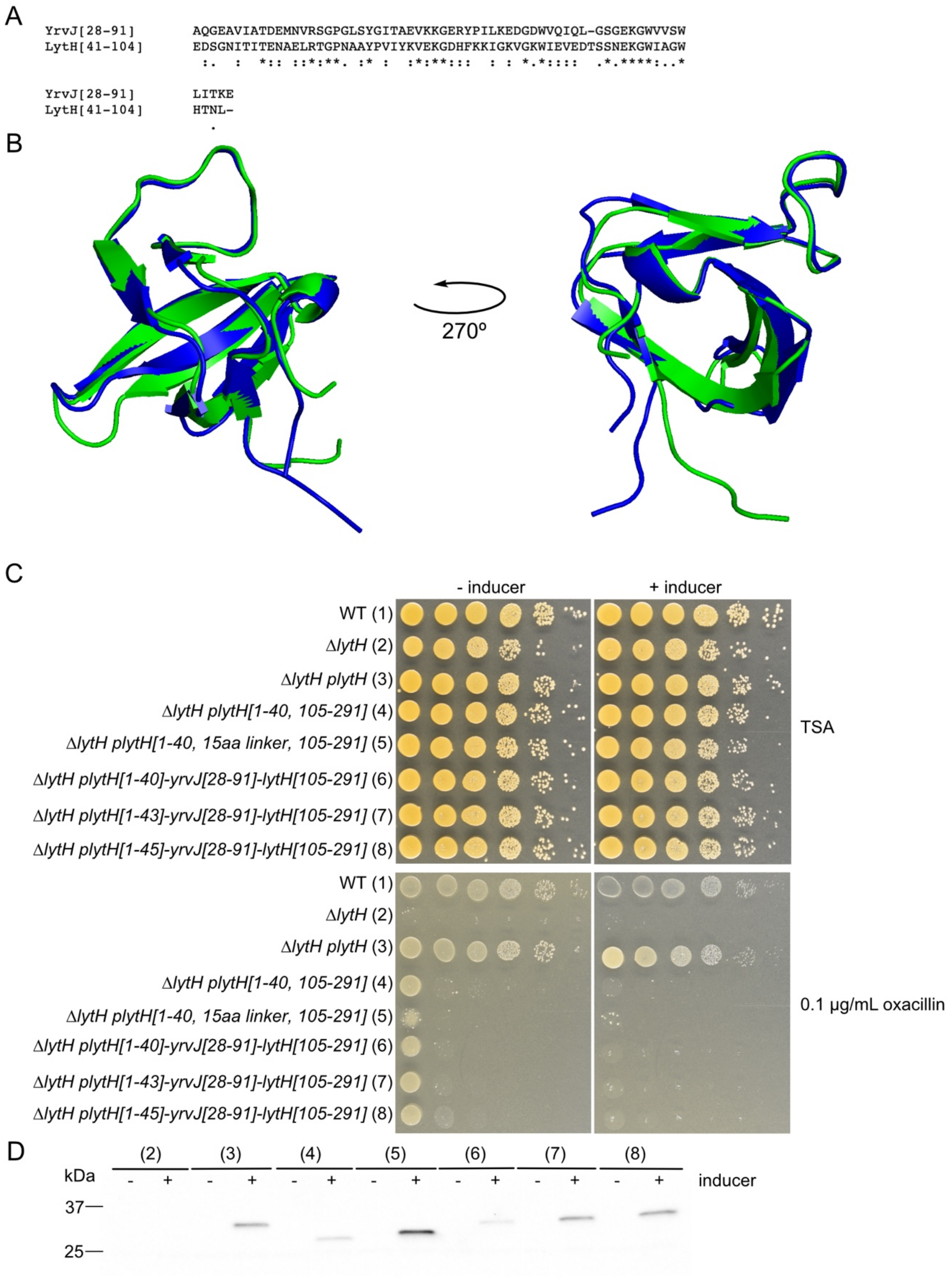


**Supplementary Figure 9. The SH3 domain of LytH is required for amidase activity in cells.** (*A*) Alignment of the sequences for the LytH and *Bacillus subtilis* amidase YrvJ SH3 domains. (*B*) Alignment of AlphaFold2-predicted structures of the LytH (green) and YrvJ (blue) SH3 domains. RMSD = 0.665.

(*C*) LytH was non-functional in cells when its SH3 domain was replaced with either a linker or the SH3 domain of YrvJ from *B. subtilis.* LytH was truncated at several points before inserting the YrvJ SH3 domain to allow for varying positionings of the replacement SH3 domain. Notably leaky expression from the pLOW plasmid allows complementation by functional LytH even in the absence of inducer. (*D*) α-FLAG Western blot of strains expressing LytH variants in an HG003 Δ*lytH* background demonstrate that the lack of complementation for SH3 domain replacement constructs is not due to lack of expression.

**Supplementary Methods**

**Plasmid construction**

*pJP62, pJP151, and pJP152.* To construct pJP62 for $His_6$-ActH[K365-K487] overexpression and purification, the gene sequence for ActH was amplified by PCR using primers oJP106 and oTD161 from *S. aureus* HG003 genomic DNA, digested with EcoRI and HindIII, and ligated into the pET28b(+) protein expression vector. To construct pJP151 and pJP152 for expression and purification of $His_6$-ActH[K365-K487, R446E] and $His_6$-ActH[K365-K487, R446A] respectively, pJP62 was linearized by PCR with primers oJP241 and oJP205 for pJP151 and oJP242 and oJP205 for pJP152 to introduce the desired mutations. The linearized DNA was phosphorylated at the 5' ends using T4 polynucleotide kinase, and the ends were ligated to produce the circular plasmids.

*pJP67, pJP73, pJP168, and pJP170.* To construct pJP67 for co-expression and purification of $His_6$-ActH[K365-K487] and LytH[T102-A291]-1x FLAG, the gene sequence for ActH was amplified by PCR using primers oJP106 and oTD161 from *S. aureus* HG003 genomic DNA, digested with EcoRI and HindIII, and ligated into pTD51 also cleaved with EcoRI and HindIII. The full length ActH gene sequence in pTD51 was thus replaced with the gene sequence for ActH[365-487]. The resulting plasmid was then linearized by PCR using primers oJP119 and oJP120. The linearized DNA was phosphorylated at the 5' ends using T4 polynucleotide kinase, and the ends were ligated to produce the circular plasmid. To construct pJP73, pJP168, and pJP170 for co-expression and purification of $His_6$-ActH[K365-K487] and LytH[T102-A291, D195A]-1x FLAG, LytH[T102-A291, H128A]-1x FLAG or LytH[T102-A291, E145A]-1x FLAG respectively, pJP67 was linearized by PCR with primers oTD103 and oTD104 for pJP73, oTD97 and oTD98 for pJP168, and oTD101 and oTD102 for pJP170 to introduce the desired mutations. The linearized DNA was phosphorylated at the 5' ends using T4 polynucleotide kinase, and the ends were ligated to produce the circular plasmids.

*pJP69 and pJP126.* To construct pJP69 for expression and purification of LytH[T102-A291]-1x FLAG, the LytH gene sequence was amplified by PCR with primers oJP122 and oTD258 and pTD78 as a template. This insert was then digested with NcoI and HindIII and ligated into pET28b(+). To construct pJP126 for expression and purification of LytH[T102-A291, D264R]-1x FLAG, pJP69 was linearized by PCR with primers oJP212 and oJP213 to introduce the desired mutation. The linearized DNA was phosphorylated at the 5' ends using T4 polynucleotide kinase, and the ends were ligated to produce the circular plasmid.

*pJP78.* To construct pJP78 for expression and purification of $His_6$-LytH[T102-A291, D195A], pTD3 was linearized by PCR with oTD103 and oTD104. The linearized DNA was phosphorylated at the 5' ends using T4 polynucleotide kinase, and the ends were ligated to produce the circular plasmid.

*pJP85 and pJP107.* To construct pJP85 for co-expression and purification of $His_6$-HRV 3C-ActH[K365-T479] and LytH[L117-A291], an insert was first amplified by PCR with primers oJP132 and oJP108 using pJP67 as the template to introduce the HRV 3C cleavage site and remove the FLAG tag. This insert was digested with EcoRI and KpnI and ligated into the pETDUET-1 backbone. The resulting plasmid was then digested with EcoRI and HindIII, and the desired gene sequence for ActH, which had been amplified by PCR using primers oJP132 and oJP133 from *S. aureus* HG003 genomic DNA and digested with EcoRI and HindIII, was ligated in. The resulting plasmid was then linearized by PCR with primers oJP119 and oJP134. The linearized DNA was phosphorylated at the 5' ends using T4 polynucleotide kinase, and the ends were ligated to produce the circular plasmid. To construct pJP107 for co-expression and purification of $His_6$-HRV 3C-ActH[K365-T479] and LytH[L117-A291, R245A], pJP85 was linearized by PCR with primers oJP173 and oJP174 to introduce the desired mutation. The linearized DNA was phosphorylated at the 5' ends using T4 polynucleotide kinase, and the ends were ligated to produce the circular plasmid.

*pJP114*. To construct pJP114 for expression of ActH-1x FLAG in *S. aureus*, the ActH gene sequence with the native actH ribosome-binding site (-25) was amplified from *S. aureus* HG003 genomic DNA by PCR with primers oJP184 and oJP185, digested with BlpI and KpnI, and ligated into pTP63.

*pJP116.* To construct pJP116 for use as an intermediary cloning plasmid, the ActH gene sequence with the native actH ribosome-binding site (-25) was amplified from *S. aureus* HG003 genomic DNA by PCR with primers oJP184 and oJP186, digested with BlpI and KpnI, and ligated into pTP63. To introduce the FLAG tag, the resulting plasmid was then linearized by PCR with primers oJP187 and oJP188. The linearized DNA was phosphorylated at the 5' ends using T4 polynucleotide kinase, and the ends were ligated to produce the circular plasmid.

*pJP129 and pJP130.* To construct pJP129 and pJP130 for expression of 1x FLAG-ActH[N2-D367] and 1x FLAG-ActH[N2-L178, K365-K487] respectively in *S. aureus*, pJP116 was linearized with primers oJP220 and oJP221 for pJP129 and oJP222 and oJP223 for pJP130 by PCR. The linearized DNA was phosphorylated at the 5' ends using T4 polynucleotide kinase, and the ends were ligated to produce the circular plasmids.

*pJP132.* To construct pJP132 for expression of ActH[K151-L178, K365-K487]-1x FLAG in *S. aureus*, pJP114 was linearized by PCR with primers oJP222 and oJP223. The linearized DNA was phosphorylated at the 5' ends using T4 polynucleotide kinase, and the ends were ligated to produce a circular plasmid. The resulting plasmid was then linearized by PCR with primers oJP225 and oJP226. The linearized DNA was phosphorylated at the 5' ends using T4 polynucleotide kinase, and the ends were ligated to produce the circular plasmid.

*pJP131 and pJP133.* To construct pJP131 as a cloning intermediate and pJP133 for expression of ActH[K337-K487]-1x FLAG in *S. aureus*, pJP114 was linearized by PCR with primers oJP222 and oJP223 for pJP131 and oJP224 and oJP225 for pJP133. The linearized DNA was phosphorylated at the 5' ends using T4 polynucleotide kinase, and the ends were ligated to produce the circular plasmids.

*pJP140-pJP144.* For each of these constructs, the gene sequence with the native rpoB ribosome-binding site (-25) was amplified by PCR. To construct pJP140 for expression of ActH-1x FLAG in *S. aureus*, the ActH gene sequence was amplified from pJP114 by PCR with primers oJP233 and oJP234. To construct pJP141 for expression of 1x FLAG-ActH[N2-D367] in *S. aureus,* the ActH gene sequence was amplified from pJP129 by PCR with primers oJP235 and oJP236. To construct pJP142 for expression of ActH[M1-L178, K365-K487]-1x FLAG in *S. aureus*, the ActH gene sequence was amplified from pJP131 by PCR with primers oJP233 and oJP234. To construct pJP143 for expression of ActH[K151-L178, K365-K487]-1x FLAG in *S. aureus,* the ActH gene sequence was amplified from pJP132 by PCR with primers oJP237 and oJP234. To construct pJP144 for expression of ActH[K337-K487]-1x FLAG in *S. aureus,* the ActH gene sequence was amplified from pJP133 by PCR with primers oJP238 and oJP234. All of these inserts were then digested with SalI and BamHI and ligated into the pLOW vector.

*pTD35*. To construct pTD35 for expression of LytH[M1-S40, D105-A291]-1x FLAG, a gBlock gene fragment with the native rpoB ribosome-binding site (-25) was synthesized by Integrated DNA Technologies (IDT). This fragment was digested with SalI and BamHI and ligated into the pLOW vector.

*pJP155.* To construct pJP155 for expression of LytH[M1-S40]-YrvJ[A28-E91]-LytH[D105-A291]-1x FLAG in *S. aureus*, pTD35 was first linearized by PCR with primers oTD95 and oTD96. The desired gene sequence for YrvJ was amplified by PCR from *Bacillus subtilis Δ6* genomic DNA with primers oJP244 and oJP245. The YrvJ insert was ligated to the linearized pTD35 DNA using the In-fusion HD Cloning Plus kit. The resulting plasmid was isolated, digested with SalI and BamHI, and the insert containing the coding sequence was ligated between the SalI and BamHI restriction sites in a fresh pLOW vector backbone.

*pJP156.* To construct pJP156 for expression of LytH[M1-S40]-$(GGS)_5$-LytH[D105-A291]-1x FLAG in *S.* aureus, pTD35 was linearized by PCR with primers oJP246 and oJP248 to insert the $(GGS)_5$ linker. The linearized DNA was phosphorylated at the 5' ends using T4 polynucleotide kinase, and the ends were ligated to produce a circular plasmid.

*pJP165 and pJP167.* To construct pJP165 and pJP167 for expression of LytH[M1-N45]-YrvJ[A28-E91]-LytH[D105-A291]-1x FLAG and LytH[M1-S43]-YrvJ[A28-E91]-LytH[D105-A291]-1x FLAG respectively in *S. aureus,* pJP155 was linearized by PCR with primers oTD95 and oJP259 for pJP165 and oTD95 and oJP260 for pJP167. The linearized DNA was phosphorylated at the 5' ends using T4 polynucleotide kinase, and the ends were ligated to produce circular plasmids. These plasmids were digested with SalI and BamHI, and the inserts containing the coding sequences were ligated between the SalI and BamHI restriction sites in a fresh pLOW vector backbone.

**Supplementary Table 2. Bacterial strains used in this study**

| Strain | Description* | Reference |
|---|---|---|
| *E. coli* | | |

| | | |
|---|---|---|
| NEB 10-beta | Host strain for plasmid cloning | New England Biolabs |
| Stellar | Host strain for plasmid cloning | Takara Clontech |
| BL21(DE3) | BL21(DE3) derivative strain for protein production | Novagen |
| C43(DE3) | BL21(DE3) derivative strain for protein production | Lucigen |
| DC10B | Δ*dcm* derivative of DH10B | (1) |
| *S. aureus* | | |
| RN4220 | Wild-type | (2) |
| HG003 | Wild-type | (3) |
| TD011 | RN4220 (pTP44), $tet^R$ | (4) |
| TD024 | HG003 Δ*lytH::*$kan^R$ | (5) |
| TD134 | HG003 Δ*lytH::*$kan^R$ (pTD30) | (5) |
| TD156 | HG003 ΔlytH::$kan^R$ (pTD35) | This study |
| TD177 | HG003 Δ*actH::*$tet^R$ | (5) |
| JP299 | HG003 Δ*actH::*$tet^R$ (pJP114) | This study |
| JP331 | HG003 Δ*actH::*$tet^R$ (pJP129) | This study |
| JP332 | HG003 Δ*actH::*$tet^R$ (pJP130) | This study |
| JP334 | HG003 Δ*actH::*$tet^R$ (pJP132) | This study |
| JP335 | HG003 Δ*actH::*$tet^R$ (pJP133) | This study |
| JP367 | HG003 Δ*actH::*$tet^R$ (pJP141) | This study |
| JP368 | HG003 Δ*actH::*$tet^R$ (pJP142) | This study |
| JP373 | HG003 Δ*actH::*$tet^R$ (pJP140) | This study |
| JP374 | HG003 Δ*actH::*$tet^R$ (pJP143) | This study |
| JP375 | HG003 Δ*actH::*$tet^R$ (pJP144) | This study |
| JP391 | HG003 ΔlytH::$kan^R$ (pJP155) | This study |
| JP392 | HG003 ΔlytH::$kan^R$ (pJP156) | This study |
| JP400 | HG003 ΔlytH::$kan^R$ (pJP165) | This study |
| JP401 | HG003 ΔlytH::$kan^R$ (pJP167) | This study |

*Abbreviations: $amp^R$, ampicillin/carbenicillin resistance; $cam^R$, chloramphenicol resistance; $erm^R$, erythromycin resistance; $kan^R$, kanamycin resistance; $tet^R$, tetracycline resistance

### Supplementary Table 3. Plasmids used in this study

| Plasmid | Description* | Reference |
|---|---|---|
| pTP63 | atc-inducible, integrative $cam^R$ expression vector for *S. aureus* | (4) |
| pLOW | IPTG-inducible, low-copy $erm^R$ expression vector for *S. aureus* | (6) |
| pET15b | $amp^R$ protein expression vector | Novagen |
| pET28b(+) | $kan^R$ protein expression vector | Novagen |
| pETDuet-1 | $amp^R$ vector for dual protein expression | EMD Millipore |
| pTD2 | pET15b-$His_6$-*lytH*[E41-A291] expression vector | (5) |
| pTD3 | pET15b-$His_6$-*lytH*[T102-A291] expression vector | (5) |
| pTD30 | pLOW-*lytH*[M1-A291]-1x FLAG containing *rpoB* RBS | (5) |
| pTD35 | pLOW-*lytH*[M1-S40, D105-A291]-1x FLAG containing *rpoB* RBS | This study |
| pTD42 | pET28b(+)-*lytH*[M1-A291]-$His_6$ expression vector | (5) |
| pTD51 | pETDUET-1-$His_6$-*actH*[N2-K487]-*lytH*[M1-A291]-1x FLAG dual expression vector | (5) |
| pTD52 | pET28b(+)-$His_6$-*actH*[N2-K487] expression vector | (5) |
| pTD78 | pET28b(+)-*lytH*[M1-A291]-1x FLAG expression vector | (5) |
| pJP62 | pET28b(+)-$His_6$-*actH*[K365-K487] expression vector | This study |
| pJP67 | pETDUET-1-$His_6$-*actH*[K365-K487]-*lytH*[T102-A291]-1x FLAG dual expression vector | This study |
| pJP69 | pET28b(+)-*lytH*[T102-A291]-1x FLAG expression vector | This study |
| pJP73 | pETDUET-1-$His_6$-*actH*[K365-K487]-*lytH*[T102-A291, D195A]-1x FLAG dual expression vector | This study |
| pJP78 | pET15b-$His_6$-*lytH*[T102-A291, D195A] expression vector | This study |

| | | |
|---|---|---|
| pJP85 | pETDUET-1-His$_6$-HRV 3C-*actH*[K365-T479]-*lytH*[L117-A291] dual expression vector | This study |
| pJP107 | pETDUET-1-His$_6$-HRV 3C-*actH*[K365-T479]-*lytH*[L117-A291, R245A] dual expression vector | This study |
| pJP114 | pTP63-*actH*[M1-K487]-1x FLAG containing *actH* RBS | This study |
| pJP116 | pTP63-1x FLAG-*actH*[N2-K487] containing *actH* RBS | This study |
| pJP126 | pET28b(+)-*lytH*[T102-A291, D264R]-1x FLAG expression vector | This study |
| pJP129 | pTP63-1x FLAG-*actH*[N2-D367] containing *actH* RBS | This study |
| pJP130 | pTP63-1x FLAG-*actH*[N2-L178, K365-K487] containing *actH* RBS | This study |
| pJP131 | pTP63-*actH*[M1-L178, K365-K487]-1x FLAG containing *actH* RBS | This study |
| pJP132 | pTP63-*actH*[K151-L178, K365-K487]-1x FLAG containing *actH* RBS | This study |
| pJP133 | pTP63-*actH*[K337-K487]-1x FLAG containing *actH* RBS | This study |
| pJP140 | pLOW-*actH*[M1-K487]-1x FLAG containing *rpoB* RBS | This study |
| pJP141 | pLOW-1x FLAG-*actH*[N2-D367] containing *rpoB* RBS | This study |
| pJP142 | pLOW-*actH*[M1-L178, K365-K487]-1x FLAG containing *rpoB* RBS | This study |
| pJP143 | pLOW-*actH*[K151-L178, K365-K487]-1x FLAG containing *rpoB* RBS | This study |
| pJP144 | pLOW-*actH*[K337-K487]-1x FLAG containing *rpoB* RBS | This study |
| pJP151 | pET28b(+)-His$_6$-*actH*[K365-K487, R446E] expression vector | This study |
| pJP152 | pET28b(+)-His$_6$-*actH*[K365-K487, R446A] expression vector | This study |
| pJP155 | pLOW-*lytH*[M1-S40]-*yrvJ*[A28-E91]-*lytH*[D105-A291]-1x FLAG containing *rpoB* RBS | This study |
| pJP156 | pLOW-*lytH*[M1-S40]-(GGS)$_5$-*lytH*[D105-A291]-1x FLAG containing *rpoB* RBS | This study |
| pJP165 | pLOW-*lytH*[M1-N45]-*yrvJ*[A28-E91]-*lytH*[D105-A291]-1x FLAG containing *rpoB* RBS | This study |
| pJP167 | pLOW-*lytH*[M1-S43]-*yrvJ*[A28-E91]-*lytH*[D105-A291]-1x FLAG containing *rpoB* RBS | This study |
| pJP168 | pETDUET-1-His$_6$-*actH*[K365-K487]-*lytH*[T102-A291, H128A]-1x FLAG dual expression vector | This study |
| pJP170 | pETDUET-1-His$_6$-*actH*[K365-K487]-*lytH*[T102-A291, E145A]-1x FLAG dual expression vector | This study |

*Abbreviations: amp$^R$, ampicillin/carbenicillin resistance; cam$^R$, chloramphenicol resistance; erm$^R$, erythromycin resistance; kan$^R$, kanamycin resistance; atc, anhydrotetracycline; IPTG, isopropyl-β-D-1-thiogalactopyranoside; HRV 3C, HRV 3C protease cleavage site; RBS, ribosome-binding site

### Supplementary Table 4. Oligonucleotide primers used in this study

| Primer | Sequence (5'-3') |
|---|---|
| oJP106 | ATCCGAATTCAAAAGAAGATAATATTTATAATAAATTGATCAAAGATGATATG |
| oJP108 | CGAGGGTACCCTACGCAGAAAAATAAATTTTAAGGCCATC |
| oJP119 | CATATGTATATCTCCTTCTTATACTTAACTAATATA |
| oJP120 | ACAAATTTAGATATTGTCGCGGATAATACG |
| oJP122 | TATACCATGGGCACAAATTTAGATATTGTCGCGGATAATACG |
| oJP132 | ATCCGAATTCATTGGAAGTTCTGTTCCAAGGCCCGAAAGAAGATAATATTTATAATAAA TTGATCAAAGATGATATG |
| oJP133 | CCGCAAGCTTTTAAGTCAACTCTTTTTCTAAGTTAATATAATCTGTAT |
| oJP134 | TTGCAAGGTAAAACAATAGTGCTTGATC |
| oJP173 | GCACAAACAAAAGTTCCTGCTGTTTTATTAGAATTA |
| oJP174 | TAACACTTGATAATTTTCTTGTCTTGAACCG |
| oJP184 | TGATGGTACCATCAAGCAGCATAGTGGAGGACTA |
| oJP185 | CGGCGCTCAGCTTACTTGTCGTCATCGTCTTTGTAGTCTTTATTTTTCGACTCATTTGATT TAGTCAACTC |
| oJP186 | CGGCGCTCAGCTTATTTATTTTTCGACTCATTTGATTTAGTCAACTC |
| oJP187 | GATGACGACAAGAACATAGACAAACAATTTTGGAAAACAATATATTATTG |
| oJP188 | GTCTTTGTAGTCCATTTAGTCCTCCACTATGCTGCTTGA |
| oJP205 | ATTTGCTATCGCTAACTCAAAATTTAATAAACC |

| | |
|---|---|
| oJP212 | GAGAAACGATGATTAAAGATCAATTACATAGACA |
| oJP213 | TAGTTGGGTTACTAATATAACCTAATTCTAATAAAAC |
| oJP220 | TAAGCTGAGCGCCGGTCGC |
| oJP221 | ATCTTCTTTAATTGTAAAAATTCTAATTTGAAGTGC |
| oJP222 | AAAGAAGATAATATTTATAATAAATTGATCAAAGATGATATGA |
| oJP223 | TAAATATAAAATCATACATAACCATATTAAGACATTTACAAA |
| oJP224 | AAAGTGAATCGTAATATTTTTGGATTTTACTAATTG |
| oJP225 | CATTTAGTCCTCCACTATGCTGCTT |
| oJP226 | AAATATATGCAGAAATTTTCACCGGCAAC |
| oJP233 | GCAGGTCGACCATAATTTTTGAGGGGTGAATCTGTATGAACATAGACAAACAATTTTGG AAAACAATATATTA |
| oJP234 | CCGGGGATCCTTACTTGTCGTCATCGTCTTTGTAGTC |
| oJP235 | GCAGGTCGACCATAATTTTTGAGGGGTGAATCTGTATGGACTACAAAGACGATGACGAC |
| oJP236 | CCGGGGATCCTTAATCTTCTTTAATTGTAAAAATTCTAATTTGAAGTGC |
| oJP237 | GCAGGTCGACCATAATTTTTGAGGGGTGAATCTGTATGAAATATATGCAGAAATTTTCA CCGGC |
| oJP238 | GCAGGTCGACCATAATTTTTGAGGGGTGAATCTGTATGAAAGTGAATCGTAATATTTTTT GGATTTTACTAAT |
| oJP241 | GAATCATTAAATGATGATGAAAAAGCATTAAAATATGTG |
| oJP242 | GCTTCATTAAATGATGATGAAAAAGCATTAAAATATGTG |
| oJP244 | TATTGCTGAATAGCAATAGTGCTCAAGGAGAAGCGGTCATC |
| oJP245 | ATCCGCGACAATATCTTCCTTTGTAATAAGCCAGGAGACTAC |
| oJP246 | CCGCTACCTCCGCTACCTCCACTATTGCTATTCAGCAATAAAAATAAAAAGATGATAA |
| oJP248 | AGGCTCTGGCGGCTCTGGCGGCTCTGATATTGTCGCGGATAATACGAAGGAGA |
| oJP259 | GAAGATAGTGGGAACGCTCAAGGAGAAGCGGTCATC |
| oJP260 | GAAGATAGTGCTCAAGGAGAAGCGGTCATC |
| oTD95 | ACTATTGCTATTCAGCAATAAAAATAAAAAGATGATAAATAAGAC |
| oTD96 | GATATTGTCGCGGATAATACGAAGGAGA |
| oTD97 | ACCAGGATCAAGCACTATTGTTTTACCTTGC |
| oTD98 | GCTGGAGGTAGTGACCAGGGTGCTTC |
| oTD101 | TAAACTTTTATATTTAGTATTGCTTGAAGCACCCTGG |
| oTD102 | GCTAAAGATTATACGTTGAAAACAGCAAAAGAATTGCAGC |
| oTD103 | ATTATGTATACTCAAATAGGCATCGCCTTTGATATC |
| oTD104 | GCTGCGTTAGAATCATCTAATGCAAATGGAATGACAG |
| oTD161 | CCGCAAGCTTTTATTTATTTTTCGACTCATTTGATTTAGTCAACTC |
| oTD258 | CCGCAAGCTTCTACTTGTCGTCATCGTCTTTGTAGT |
